# Early life adversity and adult social relationships have independent effects on survival in a wild animal model of aging

**DOI:** 10.1101/2022.09.06.506810

**Authors:** Elizabeth C. Lange, Shuxi Zeng, Fernando A. Campos, Fan Li, Jenny Tung, Elizabeth A. Archie, Susan C. Alberts

## Abstract

Does social isolation in adulthood predict survival because socially isolated individuals are already unhealthy due to adversity earlier in life (health selection)? Or do adult social environments directly cause poor health and increased mortality risk (“social causation”)? These alternative hypotheses are difficult to disentangle in humans because prospective data on survival and the environment for both early life and adulthood are rarely available. Using data from the baboon population of Amboseli, Kenya, a model for human behavior and aging, we show that early adversity and adult social isolation contribute independently to reduced adult survival, in support of both health selection and social causation. Further, strong social bonds and high social status can buffer some negative effects of early adversity on survival. These results support a growing change in perspective, away from “either-or” hypotheses and towards a multi-causal perspective that points to multiple opportunities to mitigate the effects of social adversity.

**Teaser:** Early life environments and adult social bonds have strong, but largely independent effects on survival in wild baboons.

## Introduction

In humans and other animals, the experience of harsh conditions in early life can have profound effects on adult health and survival (*1-5*). For example, one recent study found that American children who experience more than three sources of socioenvironmental adversity before age 18 can expect a 9.5 year reduction in quality-adjusted adult life expectancy (*1*). Importantly, adversity during early life is also linked to social adversity in adulthood, including both low socioeconomic status (SES) and challenges in forming strong and supportive social relationships (*6-8*). In turn, low SES and social isolation/low social support are linked to poor health and all-cause mortality (*9-12*). However, while early life adversity, poor adult social relationships, and low adult social status have all been linked to poor adult survival, the causal relationships between these factors are not well understood.

Specifically, while most hypotheses acknowledge that early life environments can affect both adult health and the adult social environment (Figure 1; *13, 14*, competing hypotheses differ in the extent to which they identify adult health (driven by early adversity) as the cause or consequence of differences in adult social relationships. For instance, the health selection hypothesis posits that poor health status affects the adult social environment, preventing attainment of high social (or socioeconomic) status and compromising the formation of strong social relationships (*15*; Figure 1A). Under this scenario, poor health—arising from early life adversity or some other source—is the primary cause of both adverse social environments and poor health/survival in adulthood. Alternatively, the social causation hypothesis posits that social isolation and low social status in adulthood play a direct, causal role in the connection between the social environment, health, and survival (*10, 16*). Under this scenario, poor social relationships and/or low social status in adulthood are sufficient to trigger poor health/survival in adulthood (Figure 1B). Thus, while early life adversity may also contribute to variation in health and/or the social environment, enhancements to the social environment in adulthood are viable paths to improving adult health and lifespan. The causal effects of adult social environments may act in parallel to the effects of early adversity or function as a source of resilience against the costs of early adversity (i.e., the “social buffering” hypothesis; *17-19*).

**Figure 1.**
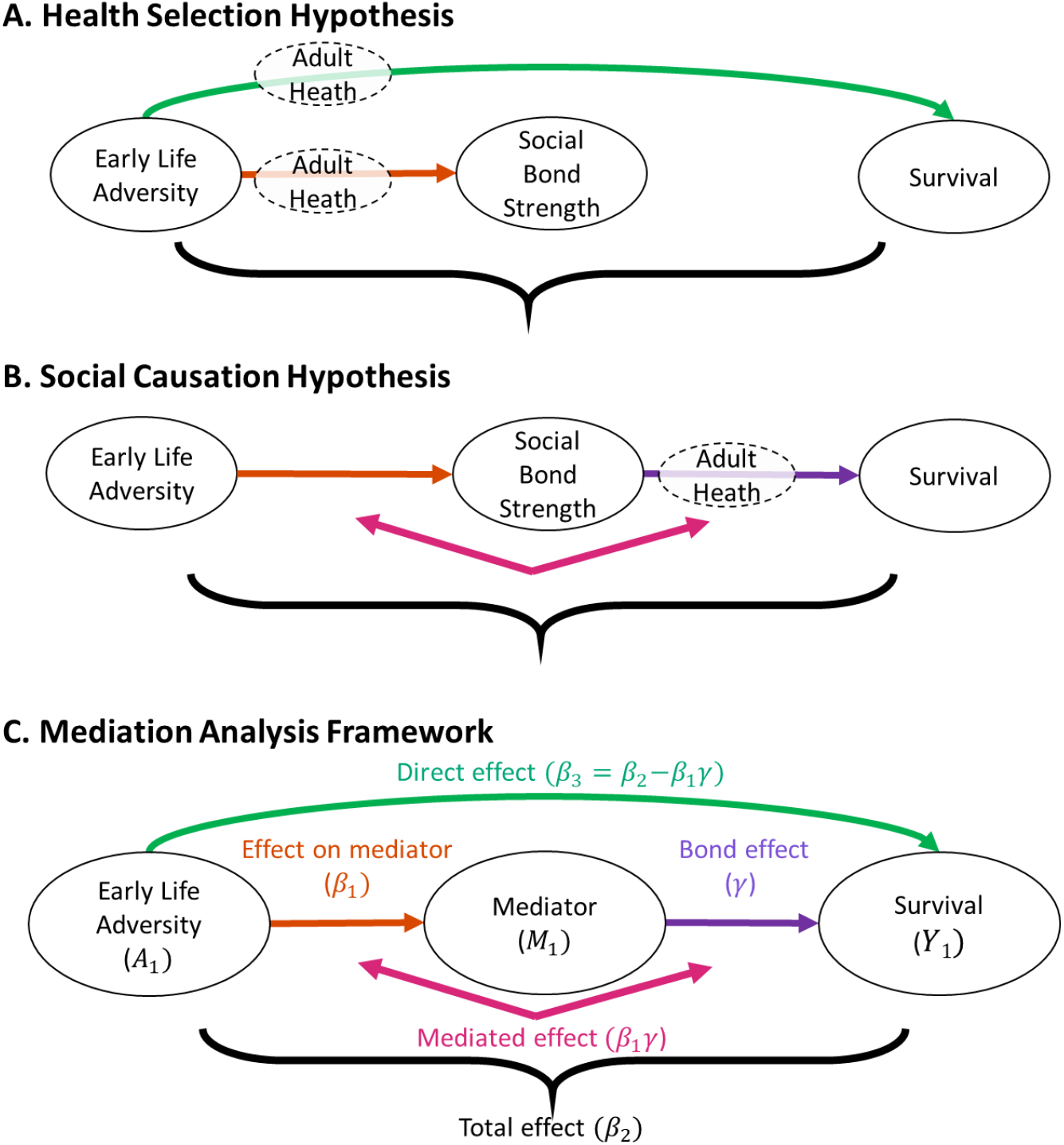
Hypotheses and mediation analysis framework and hypotheses linking early life adversity, adult social bond strength, and survival. The health selection hypothesis (A) posits that poor adult health arising from early life adversity prevents individuals from forming strong social relationships. Under health selection, we predict a link between early adversity and adult social bond strength (orange arrow), and a direct link between early adversity and survival (green arrow) outside of the pathway that includes social bond strength, but no mediated effect (pink arrow in B, C), and no independent effects of social bond strength on survival (purple arrow in B, C). The social causation hypothesis (B) predicts that social bond strength is a direct cause of survival differences (purple arrow). It also predicts that any effects of early life environments on survival, at least as they relate to social relationships, are due to a mediated effect (pink arrow) where early adversity affects adult social bond strength (orange arrow), which in turn affects survival (purple arrow). Under social causation, early life adversity may affect survival via other pathways (e.g., green arrow in A), but social relationships have an important causal effect. (C) The mediation analysis models the links between early life adversity (*A*_1_), adult mediator phenotypes (*M*_*1*_, social bond strength with females or males), and survival (*Y*_1_). Mediation models produce estimates of (i) the direct effect of early life adversity on survival outside of the pathway that includes the mediator (*β*_3_, green arrow), (ii) the mediated effect of early life adversity on survival through the pathway that includes the mediator (*β*_1_*γ*, pink arrow), (iii) the effect of early life adversity on the mediator (*β*_1_, orange arrow), (iv) the effect of the mediator on survival independent of early adversity (*γ*, purple arrow, the bond effect), and (v) the total effects on survival (*β*_2_, black bracket). Note that the expressions *β*_1_*γ* and (*β*_3_ = *β*_2_ − *β*_1_*γ*) in panel C hold exactly only when all models (between *A, M*, and *Y*) are linear. Here we use these merely as notations (instead of mathematical equations) to label the qualitative relationship between total, mediated, direct, and bond effects.

Distinguishing between health selection and social causation is important to both evolutionary biologists and social scientists. Understanding what forces drive variation in survival helps to identify the traits targeted by natural selection, shedding light on the evolutionary underpinnings of early life effects and sociality. At the same time, understanding the causes of variation in health and mortality can inform investment in public health interventions and policy. However, despite extensive research on the pathways linking early experience and adult life outcomes, the relative importance of health selection versus social causation in explaining social environmental effects on adult survival is widely debated (*13, 20*-*24*).

To address this debate, the best approach is to link prospectively collected data on early life adversity and prospectively collected information on the adult social environment and survival in the same individuals (*20, 25*). Existing data typically do not permit such analyses in human populations, but appropriate data are sometimes available for wild animal populations that have been under continuous observation for many years (*9*). Further, the social determinants of health in many social mammal species resemble those described in humans, making wild animal models a useful tool for dissecting the relationships among early life adversity, adult social behavior, and lifespan. For example, in several nonhuman mammals, early life adversity is linked to low adult social status or weak adult social relationships (*26-29*). Similarly, low social status or weak social relationships are associated with higher mortality rates in a range of social mammal species (*9*).

In this study, we use a mediation analysis framework to examine the relationships among early life adversity, adult social behavior, and survival in an established wild animal model of aging: the baboons studied by the Amboseli Baboon Research Project in the Amboseli ecosystem, Kenya (*30, 31*). Our goals were to determine the relative importance of health selection and social causation in explaining survival patterns in adult female baboons, and to determine whether adult social relationships buffer the effects of early life adversity. We focused on adult females because male baboons disperse from their natal social groups when they mature, making it difficult to distinguish male dispersal from death (*32*).

If health selection explains the link between adult phenotypes and survival, two predictions ensue: (i) early adversity should predict weak adult social relationships, and (ii) early adversity should have a direct effect on adult survival that is not mediated by adult social relationships (Figure 1A). This pattern would support the idea that poor health arising from early adversity leads to both social isolation and poor survival in adulthood. In contrast, social causation predicts (i) that adult social bonds have strong, direct effects on survival, and (ii) that these effects occur regardless of a female’s early life experience (Figure 1B). Such a pattern would support the idea that the effect of early life adversity on survival is at least partly mediated by its effects on adult social relationships. Finally, the social buffering hypothesis, which is consistent with social causation, would be supported if the effects of early life adversity on adult survival are moderated by adult social behavior.

Previous work on female baboons in Amboseli has shown that harsh early life environments predict reduced adult female lifespan (*26, 33*) as well as a moderate degree of social isolation in adulthood (*26, 34*). In addition, in adult females, weak social bonds predict decreased lifespan (*35, 36*). However, no previous study in either animals or humans has sought to prospectively link early life adversity, adult social behavior, and survival in an integrated analysis. Therefore, it is unknown if adult social relationships have independent effects on survival or merely act as mediators that link early life to survival. We focus on adult female social bonds as candidate mediators, and we exclude adult social status as a potential mediator for two reasons: (i) previous studies in this population find no effects of female social status on survival (*35, 36*), and (ii) preliminary analyses using our mediation framework ruled out social status as a potential mediator of early life adversity and demonstrated that social status is not influenced by cumulative early life adversity (see Materials and Methods). However, in our test of the social buffering hypothesis, we consider both adult social bonds and adult social status as possible moderators of the relationship between early life and survival.

## Mediation and moderation frameworks

### Mediation models

Our mediation analysis framework is based on structural equation models that examine the links between early life, adult social phenotypes, and survival (Figure 1C, *37-39*). The 199 females in this study were observed from birth and survived to at least four years old, approximately the earliest age of reproductive maturation (average age at menarche = 4.73 ± 0.56 years). For each female, we evaluated her exposure to six different adverse socioenvironmental conditions in early life: 1) drought in the first year of life, 2) large group size at birth, 3) low maternal social status at birth, 4) low maternal social connectedness during the first two years of life, 5) the presence of a close-in-age younger sibling, and 6) maternal loss before four years of age (Table 1; *26, 33, 34*).

**Table 1.** Sources of early life adversity and the number of females that experienced each source.

| Source of Adversity | Description | N of females who experienced adversity | N of females who did not experience adversity |
| --- | --- | --- | --- |
| Drought | <200 mm of rainfall during the first year of life | 28 | 171 |
| Large group size | Group size in the top quartile (> 33 adults) at the subject's birth, indicating high social density | 32 | 167 |
| Close-in-age younger sibling | Younger sibling born less than 1.5 years after the subject's birth | 40 | 159 |
| Maternal loss | Mother dies during the first four years of the subject's life | 38 | 161 |
| Low maternal social status | Mother's proportional dominance rank in the lowest quartile at the subject's birth | 46 | 153 |
| Low maternal social connectedness | Mother's social connectedness in the lowest quartile during the subject's first two years of life | 54 | 145 |

We constructed two sets of mediation models (see Materials and Methods), each with a different mediator variable (*M*), linking the treatment (early life adversity, *A*) to survival (*Y*, measured by the hazard ratio, *λ*; Figure 1C). The two mediators we examined were quantitative measures of social bond strength with other adult females and with adult males (see *Potential mediators*, below). Because both of these variables are known to be linked to adult survival (*36*), either could act as a mediator of early life adversity. We considered a female’s social bonds with other adult females separately from her social bonds with adult males because same-sex and opposite sex social relationships have different relationships with early adversity and with survival and are not well-correlated (*26, 34-36*).

The mediation analysis enables us to break down the total effect of early life adversity on survival (*β*_2_, black arrow in Figure 1C) into direct (*β*_3_) and mediated (*β*_1_*γ*) effects. The direct effect (*β*_3_) of early life adversity on survival is the pathway connecting these variables independent of the mediator (green arrow in Figure 1C). The mediated (or indirect) effect (*β*_1_*γ*) is the pathway connecting early life adversity and survival that runs through the mediator variable; in our case, measures of social bond strength (pink arrows in Figure 1C). The mediation framework also assesses the effect of early adversity on the mediator (*β*_1_, orange arrow in Figure 1C) and the effect of the mediator on survival independent of early adversity, hereafter the ‘bond effect’ (*γ*, purple arrow in Figure 1C).

For each of our mediators, we estimated the links between early life adversity (*A*), social bond strength (*M*), and survival (*Y*, measured by the hazard ratio *λ*) by fitting three equations as proposed by Zeng, Lange, Archie, Campos, Alberts and Li (*40*); for more details see Materials and Methods). The first equation evaluates the effect of early life adversity on observed values for the mediator, conditional on covariates, *C*, and random effects, *r* (orange arrow in Figure 1C):

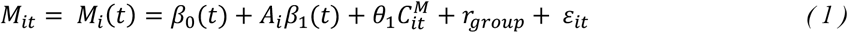

where *i* is individual and *t* is age class. Here *β*_1_represents the effect of early adversity on social bond strength. The second equation models the total effect of early life adversity on survival (e.g., the change in hazard rate related to early adversity; *β*_2_, black arrow in Figure 1C), which does not differentiate between direct and mediated effects:

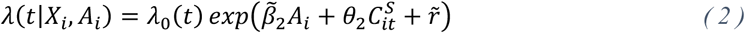

The third equation is similar to Equation 2, but incorporates estimates of the mediator based on the parameters previously fit for Equation 1. It allows us to estimate the value of the effect of the mediator on survival given the estimate of the mediator f{α, *M*_*i*_(*t*)}:

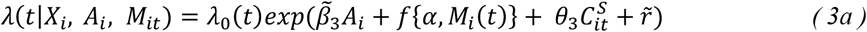

where the mediator component f{α, *M*_*i*_(*t*)} equals:

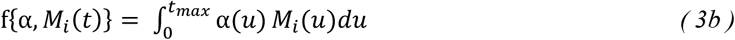

where *t*_*max*_is the maximum lagged time (here three years) and α(*u*) is a time varying constant. Equation 3b estimates the mediator for the previous three years of life, based on values for the covariates, early life adversity, and the effect sizes estimated in Equation 1 (i.e., Equation 3b is fit based on estimated values of the mediator, not directly on observed data). We designate this value the ‘three-year mediator value’, where each year corresponds to a female age class, starting on her birthday and ending one day before her subsequent birthday. We also considered models where the mediator was estimated based on the same year of life as survival (‘one-year mediator models’) and results are consistent with three-year mediator models (Tables S1, S2).

Note that the effects *β*_2_ and *β*_3_ in Figure 1C are not numerically identical to the coefficients 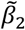 and 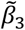 in Equation 2 and 3, respectively. While they are analogous to *β*_2_ and *β*_3_ in terms of the effects they represent, they differ because of the nonlinear hazard scale and complex functional model adopted in the analysis (i.e., in practice, we analyze a decomposition of a functional form fit to the social relationship data rather than the estimated social bond values directly; see Zeng, Lange, Archie, Campos, Alberts and Li (*40*)). Similary, the bond effect *γ* does not directly correspond to a specific model parameter. Instead, *β*_2,_ *β*_3,_ and *γ* are calculated from functions involving all parameters in Equations 1, 2 and 3 (see the Materials and Methods and derivations in Zeng, Lange, Archie, Campos, Alberts and Li (*40*)).

First, we modeled the effects of cumulative early adversity on both mediators (social bond strength with females and social bond strength with males) and on survival. We measured cumulative early adversity as a continuous variable representing the sum of the six individual sources of adversity for each subject. No individual had a cumulative adversity score greater than four (mean=1.196±0.936 SD). Second, we built multivariate models to assess the effect of each individual source of adversity on each mediator and on survival, while holding the other sources of adversity at zero. In these models of individual sources of adversity, each measure of adversity was modeled as a categorical variable (a value of one for subjects that experience the adverse event, and zero for those that did not).

### Moderation models

To test the social buffering hypothesis, which posits that adult social relationships act as a source of resilience in the face of early adversity, we next treated three adult social phenotypes (social bond strength with females, social bond strength with males, and social status) as potential moderators instead of mediators of early life adversity. In contrast to the mediation models, the moderation models test whether the social phenotypes influence the strength and direction of the effect of early life adversity on survival without making causal assumptions about the pathways involved. Moderation is captured by the interaction between the exposure *A*_*i*_ and mediator *M*_*i*_(*t*) with the interaction term A_i_g{η, M_i_(t)} in the following model:

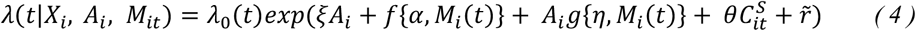

Therefore, this approach allows us to estimate how the effects of early adversity on survival vary across different levels of the social bonds or social status.

## Potential mediators and moderators

### Mediators

We measured each female’s social bond strength with females – i.e., the strength of her social bonds with her top three female partners in each year of her life – and each female’s social bond strength with males – the strength of her social bonds with her top three male partners in each year of her life – as two distinct potential mediators (*M*) of the effects of early life adversity on survival. We used grooming relationships to assess social bond strength because grooming is the most prominent affiliative behavior in baboons and many other primates (*41-44*). These mediators were represented in Equations 2 and 3 as estimates over three-year periods (Equation 3b), based on the values of their covariates and the parameters fit in Equation 1. We also estimated mediators over shorter, one-year periods, as reported in Tables S1-S2; because all analyses based on shorter periods produced qualitatively similar results, we focus on the three-year estimates here.

### Moderators

We considered adult social bond strength with females, adult bond strength with males, and adult social status as potential moderators. We assessed social bond strength using the same method described above (based on grooming relationships calculated as trajectories as in Equation 1). We assessed social status using observations of wins and losses in dyadic agonistic interactions between adult female study subjects. A female dominance matrix was created for each month based on these win/loss outcomes, and female ordinal dominance ranks were assigned by minimizing entries below the diagonal (*45*). We then scaled these ordinal rankings by group size and assigned to each female a ‘proportional dominance rank’ (*46*), calculated as [1 – (ordinal rank – 1) / (number adult females – 1)]. A female’s proportional dominance rank represents the proportion of adult females that she dominates. We first calculated annual mean values for social status for each subject, and then estimated their social status trajectories over three-year periods, given covariates and parameter estimates for an analogue of Equation 1, with *M*_*it*_ is redefined as annual mean proportional dominance rank instead of annual social bond strength (see also Methods).

## Results

### Cumulative early adversity and survival: Mediated effects are weak, direct effects are strong

As expected, we found a strong total effect (*β*_2_) of cumulative early adversity on adult female survival, recapitulating previous work (Tables 2-3; black bracket in Figure 2; black points and lines in Figures S1-S2; *26, 33*). Approximately 90% of the total effect (1.43 of 1.60 years of lost life per additional exposure and 1.45 of 1.59 years, for the models considering social bond strength with females and social bond strength with males, respectively) was explained by the direct effect (*β*_3_) of cumulative early adversity on survival, outside of the pathways that included social bonds with either sex (Tables 2-3; green arrows in Figure 2; green points and lines in Figures S1-S2). Thus, the lives of females who experience four sources of early life adversity are predicted to be 6.4 years shorter than those of females that experience none, on average. Of these six years, ∼5.6 years would be explained by the effects of early adversity on survival, independent of mediation by social bonds. Results were similar if we estimated mediation effects over shorter, one-year periods instead of three-year periods (Tables S1-S2).

**Table 2.** Mediation results from models in which social bond strength with females was the mediator. Total, direct, mediated and bond effects are measured in years. The effect on the mediator is measured in social bond strength units (i.e., DSI units; 1 SD in social bonds with females=0.59 DSI units). Bolded effects are those for which the 95% credible intervals (shown in brackets below each effect size estimate) did not overlap zero. Effect names are colored as in Figure 1.

| | Total effect<br>($\beta_2$, years) | Direct effect<br>($\beta_3$, years) | Mediated effect<br>($\beta_1\gamma$, years) | Effect on<br>mediator<br>($\beta_1$, DSI units) | Bond effect<br>($\gamma$, years) |
| --- | --- | --- | --- | --- | --- |
| Drought | **-2.70**<br>[-4.96, -0.44] | **-2.26**<br>[-4.04, -0.48] | **0.44**<br>[0.03, 0.85] | **-0.21**<br>[-0.38, -0.03] | **2.19**<br>[0.56, 3.82] |
| Large group size | -1.60<br>[-4.02, 0.83] | -1.38<br>[-2.89, 0.13] | 0.22<br>[-0.01, 0.44] | -0.11<br>[-0.22, 0.01] | **2.39**<br>[0.61, 4.17] |
| Close-in-age younger sibling | -0.90<br>[-5.45, 3.65] | -0.59<br>[-1.99, 0.81] | 0.31<br>[-0.04, 0.66] | **-0.15**<br>[-0.28, -0.03] | **2.29**<br>[0.75, 3.83] |
| Maternal loss | **-3.30**<br>[-5.79, -0.81] | **-2.67**<br>[-4.77, -0.57] | **0.63**<br>[0.06, 1.20] | **-0.26**<br>[-0.47, -0.04] | **2.60**<br>[0.79, 4.40] |
| Low maternal social connectedness | 0.10<br>[-2.07, 2.27] | 0.15<br>[-1.31, 1.62] | 0.05<br>[-0.17, 0.27] | -0.05<br>[-0.15, 0.06] | **2.60**<br>[0.87, 4.34] |
| Low maternal social status | -1.80<br>[-4.37, 0.78] | -1.36<br>[-2.90, 0.19] | 0.44<br>[-0.05, 0.93] | **-0.14**<br>[-0.25, -0.03] | **2.49**<br>[0.66, 4.32] |
| Cumulative adversity | **-1.60**<br>[-2.84, -0.36] | **-1.43**<br>[-2.52, -0.35] | **0.17**<br>[0.01, 0.32] | **-0.09**<br>[-0.16, -0.01] | **2.20**<br>[0.74, 3.65] |

**Figure 2.**
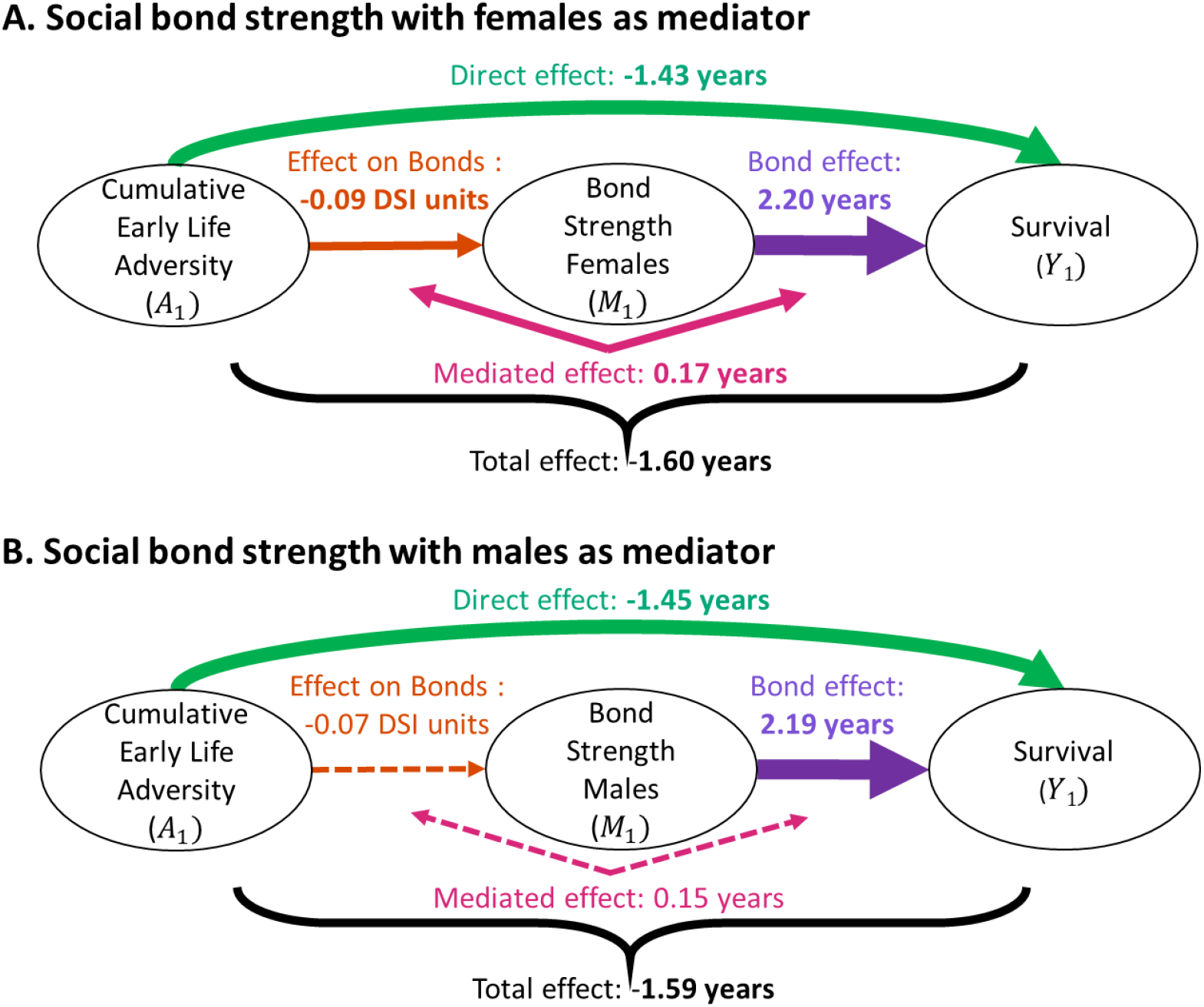
Mediation analysis results. (A) Results from our mediation model using social bond strength with adult females as the mediator. (B) Results from our mediation model using social bond strength with adult males as the mediator. Solid lines indicate effects for which 95% credible interval did not overlap zero, dashed lines indicate effects for which 95% credible interval did overlap zero.

We also found substantial effects of both mediators (*γ*) on survival, independent of effects of early life adversity. A one unit increase in social bond strength with either adult females or adult males predicted a 2.2-year improvement in survival, independent of the effects of early adversity, where one unit represents approximately 1.7 standard deviations for social bond strength with females and 1.4 SD for social bond strength with males (Tables 2-3; purple arrows in Figures 1; Figures S1-S2; see Tables S1-S2 for results with mediators estimated over shorter, one-year periods). While the effects of social bonds on survival broadly recapitulate previous findings in this population (*35, 36*), this analysis is the first to demonstrate that these effects remain strong after controlling for levels of early adversity.

Notably, despite the fact that cumulative early adversity significantly predicted weaker social bonds with females (*β*_1_, orange arrows in Figure 1C), and that stronger social bonds with both sexes predicted higher survival, mediated effects were weak in all of our models of cumulative adversity. Specifically, the pathway through social bonds with females improved lifespan by only 2.04 months (10.6%), compared to the 1.60 year reduction in lifespan for each additional source of adversity (the mediated effect, *β*_1_*γ*, pink bracket in Figures 2A, S1; Table 2). This result may stem from the fact that the effect of cumulative early adversity on social bonds, while detectable, is relatively weak: early adversity is associated with a 0.09 unit decrease in social bonds with females, which is small compared to the 1 unit increase in social bonds with females necessary to produce a 2.2 year improvement in lifespan via the bond effect. Social bonds with males did not detectably mediate the relationship between cumulative early adversity and survival (Figures 2B, S2; Table 3).

**Table 3.** Mediation results from models in which social bond strength with males was the mediator. Total, direct, mediated and bond effect are measured in years. The effect on the mediator is measured in social bond strength units (i.e., DSI units; 1 SD in social bond strength with males=0.70 DSI units). Bolded effects are those where the 95% credible intervals did not overlap zero. Effect names are colored as in Figure 1.

| | Total effect<br>($\beta_2$, years) | Direct effect<br>($\beta_3$, years) | Mediated effect<br>($\beta_1\gamma$, years) | Effect on<br>mediator<br>($\beta_1$, DSI units) | Bond effect<br>($\gamma$, years) |
| --- | --- | --- | --- | --- | --- |
| Drought | **-2.70**<br>[-4.96, -0.44] | **-2.33**<br>[-4.17, -0.50] | **0.37**<br>[0.02, 0.71] | **-0.16**<br>[-0.29, -0.02] | **2.40**<br>[0.62, 4.17] |
| Large group size | -1.60<br>[-4.01, 0.82] | -1.53<br>[-3.20, 0.15] | 0.07<br>[-1.93, 2.07] | -0.04<br>[-0.92, 0.84] | **2.39**<br>[0.61, 4.17] |
| Close-in-age younger sibling | -0.89<br>[-5.33, 3.55] | -0.69<br>[-2.08, 0.70] | 0.20<br>[-1.20, 1.60] | -0.11<br>[-0.61, 0.39] | **2.29**<br>[0.75, 3.84] |
| Maternal loss | **-3.30**<br>[-5.78, -0.81] | **-3.21**<br>[-5.73, -0.68] | 0.09<br>[-2.38, 2.56] | -0.06<br>[-1.95, 1.84] | **2.20**<br>[0.67, 3.73] |
| Low maternal social connectedness | 0.11<br>[-1.90, 2.12] | 0.37<br>[-1.26, 1.99] | 0.26<br>[-0.15, 0.66] | -0.15<br>[-0.32, 0.02] | **2.20**<br>[0.73, 3.66] |
| Low maternal social status | -1.80<br>[-4.38, 0.78] | -1.16<br>[-2.70, 0.38] | 0.64<br>[-0.07, 1.35] | **-0.25**<br>[-0.44, -0.06] | **2.30**<br>[0.61, 3.99] |
| Cumulative adversity | **-1.59**<br>[-2.82, -0.36] | **-1.45**<br>[-2.54, -0.35] | 0.15<br>[-0.33, 0.62] | -0.07<br>[-0.37, 0.22] | **2.19**<br>[0.74, 3.64] |

Taken together, our results are not fully explained by either health selection or social causation. Health selection would predict a direct effect of early adversity (presumed to compromise adult health) on both social bond strength and survival, without a strong bond effect on survival. Instead, early life adversity and social bonds both appear to have direct, independent effects on survival that are of similar magnitidues. Consequently, a female baboon who experienced higher than average (1 SD above the mean) cumulative early life adversity, adult social bond strength with females, and adult social bond strength with males would be predicted to experience a 1.35 year reduction in lifespan attributable to her early life environment, a 1.29 year improvement in lifespan attributable to her social bonds females in adulthood, and a 1.29 year improvement in lifespan attributable to her social bonds with males in adulthood. In other words, both early adversity (likely via a route through poor adult health), and adult social behavior are important in determing survival in adulthood.

We next considered whether the weak mediation we observed – in spite of effects of early adversity on social bonds and of social bonds on survival – might result from a mismatch in the timing of these effects. To explore this possibility, we designed a simulation analysis in which we defined two stages corresponding to early and late adulthood. We then assigned early life effects on the mediator, and mediator effects on survival, in all possible combinations of early and late timing of effects (see Supplementary Text: “Simulation to explore the small mediated effect”). In our simulations, we fixed the values of both the effect of early adversity on the mediator (“Effect on mediator”, orange arrow in Figure 1C) and the effect of the mediator on survival (“Isolation effect”, purple arrow in Figure 1C). Even though the component parts of the mediated effect were kept constant in the simulations, the estimate of the overall mediated effect (pink arrows in Figure 1C) depended on the timing of these effects. Mediated effects were largest when the timing of early life and mediator effects were matched; i.e., when either (i) early adversity had its strongest effects on the mediator early in life *and* the mediator had its strongest effects on survival early in life, or (ii) early adversity had its strongest effects on the mediator late in life *and* the mediator had its strongest effects on survival late in life (see Supplementary Text: “Simulation to explore the small mediated effect”, Figure S4). The results of this simulation support the idea that the timing of these effects during the life course could play a role in determining the strength of the mediated effect. They further suggest that, in the Amboseli baboons, the timing of early life effects on adult social isolation may be mismatched with the timing of social bond effects on survival. This topic merits future exploration.

### Social bonds do not mediate the effects of individual sources of early adversity

Similar to the effects of cumulative adversity, individual sources of adversity acted outside of the pathway that includes social bonds, with little evidence for mediated effects in our three-year mediator models (Tables 2-3; Figures S1-S2). More than 81% of the effects of individual sources of adversity were attributable to direct effects (87% if only considering significant direct effects). Among the six individual sources of early adversity, maternal loss and drought exerted the strongest and most consistent effects on both adult female survival and social bond strength with adult females (Tables 2-3; Figures S1-S2). Drought, but not maternal loss, was also linked to weaker social bonds with adult males. In contrast to the effects of maternal loss on social isolation from adult females, maternal loss did not predict social bond strength with adult males: the estimated effect size was near zero (0.06 DSI units; Table 3). Consistent with our main results, the effects of individual sources of adversity on survival were also not detectably mediated by measures of social bonds with either sex based on one-year intervals (Table S1-S2).

### Moderating effects: Social bonds buffer the effects of some sources of early adversity

Neither social status nor social bond strength with either sex moderated the link between cumulative adversity and survival (Table 4; Figure 3A, B; results were similar when we used moderator trajectories estimated over a shorter, one-year period, Table S3). However, social bond strength with males and social bond strength with females both moderated the link between one individual source of adversity – maternal loss – and survival. Specifically, stronger social bonds with either females or with males during adulthood buffered the negative effect of maternal loss on survival (and conversely weaker social bonds amplified the negative effect of maternal loss on survival; Table 4, Table S3: Figure 3A,B,D,E). In other words, survival was disproportionately lower for females who lost their mother early in life and were more socially isolated in adulthood (and conversely, survival was disproportionately higher for females who lost their mother but formed strong social relationships in adulthood, with either sex; Figure 3A,B,D,E). Females who lost their mother early in life but maintained strong social relationships with other females (1 SD above the mean) experienced a 10% reduction in hazard ratios relative to females who lost their mothers and had average social bond strength to other females. In contrast, females who lost their mothers and had weak social relationships with females (1 SD below the mean) had 16% higher hazard ratios than females who lost their mothers and had average social bond strength to other females (Figure 3D). The effect was stronger for bonds with males, where females who lost their mothers in early life but maintained strong social bonds with males (1 SD above the mean) had an 18% lower hazard ratio, while those who had weak social bonds with males (1 SD below the mean) had a 16% higher hazard ratio, compared to the effects of maternal loss for females with average social bond strength (Figure 3E). In addition, another individual source of early adversity – low maternal social connectedness – was buffered by strong adult social bonds with males, but not by adult social bonds with females (Figure 3B).

**Table 4.** Moderation results from models in which social bond strength with females, social bond strength with males, and female social status were the moderators. Values represent the magnitude of the interaction effects measured in log hazard ratio (HR). Bolded effects (those for which the 95% CI did not overlap zero) show that the effects of maternal loss on survival were moderated by all three phenotypes and that the effects of low maternal social connectedness were moderated by adult social relationships with males and female social status. A negative interaction effect indicates that increased adult social bond strength or higher social status acts as a buffer to reduce the negative effects of early adversity on survival. A positive interaction effect value means that adult social bond strength or higher social status acts as an amplifier to increase the negative effects of early adversity on survival.

|  | Social Bonds<br>w/ Females<br>(log HR) | Social Bonds<br>w/ Males<br>(log HR) | Social Status<br>(log HR) |
| --- | --- | --- | --- |
| Drought | -0.19<br>[-0.41, 0.03] | -0.13<br>[-0.33, 0.06] | -0.07<br>[-0.29, 0.15] |
| Large group size | 0.19<br>[-0.06, 0.44] | -0.12<br>[-0.27, 0.02] | -0.19<br>[-0.40, 0.02] |
| Close-in-age younger sibling | 0.13<br>[-0.38, 0.64] | 0.00<br>[-0.33, 0.33] | 0.25<br>[-0.19, 0.69] |
| Maternal loss | **-0.26**<br>**[-0.39, -0.13]** | **-0.21**<br>**[-0.3, -0.12]** | **-0.18**<br>**[-0.33, -0.04]** |
| Low maternal social connectedness | 0.11<br>[-0.03, 0.26] | **-0.15**<br>**[-0.24, -0.06]** | **-0.19**<br>**[-0.37, -0.01]** |
| Low maternal social status | 0.02<br>[-0.15, 0.18] | 0.04<br>[-0.04, 0.13] | -0.02<br>[-0.19, 0.16] |
| Cumulative adversity | -0.02<br>[-0.05, 0.01] | -0.03<br>[-0.05, 0.00] | 0.02<br>[-0.02, 0.06] |

**Figure 3.**
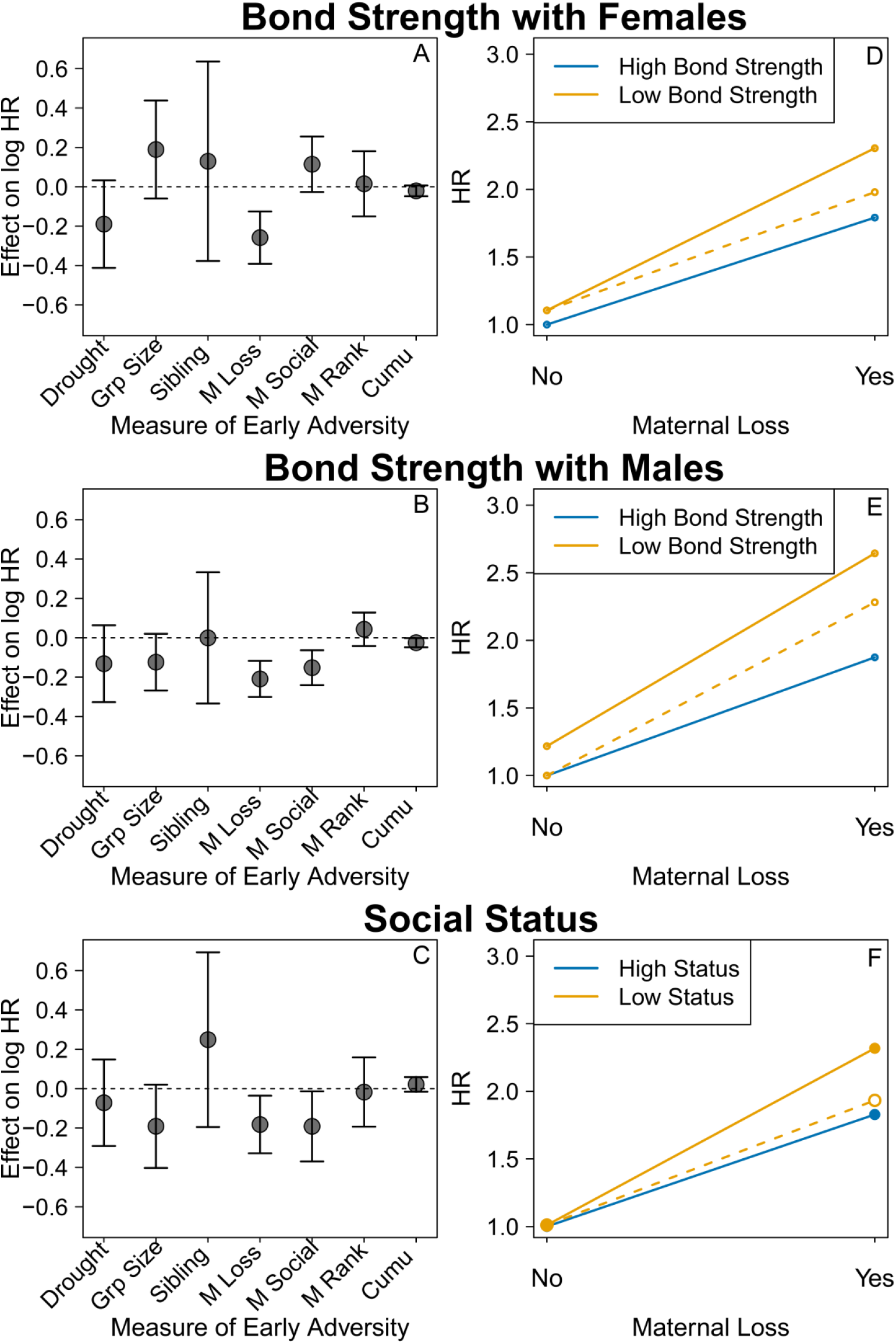
Moderation models support some forms of social buffering. (A, B, and C) Moderating effects of (A) social bond strength with females, (B) social bond strength with males, and (C) social status on the relationship between early adversity and survival (log of the hazard ratio, HR). A positive value on the y-axis means that greater social bond strength or higher social status amplify the negative effects of early adversity on survival. Grp Size indicates large social group size, M Loss indicates maternal loss, M Social indicates low maternal social connectedness, M Rank indicates low maternal rank, and Cumu indicates cumulative adversity. Panel A shows that strong social bonds with females buffer the effects of maternal loss; Panel B shows that strong social bonds with males buffer the effects of both maternal loss and low maternal social connectedness; Panel C shows that high social status buffers the effects of both maternal loss and low maternal social connectedness. (D, E, and, F) The effects of (D) social bond strength with females, (E) social bond strength with males, and (F) social status on the relationship between maternal loss and survival (measured as the hazard ratio, HR). The orange dashed line in each panel represents the expected effect of maternal loss on the hazard ratio for adult females in the absence of any moderating effects of social bonds or status. The blue solid line shows that females with social bond strength one standard deviation (SD) above the mean (i.e., females with stronger social bonds) or females with social status one SD above the mean (i.e., females with high status) experience a disproportionately lower hazard ratio in the presence of maternal loss. The orange solid line shows that females with social bond strength or social status one SD below the mean experience a disproportionately higher hazard ratio as a function of maternal loss.

Female social status also moderated early life maternal loss and low maternal social connectedness effects on survival (Figure 3C, Table 4; note that this effect was not detectable when moderator trajectories were estimated over a shorter, one-year period, Table S3). Specifically, survival was disproportionally lower for low-ranking females who lost their mothers early in life or had a socially isolated mother, and disproportionally higher for high-ranking females who lost their mothers early in life or had a socially isolated mother (Figure 3C,F). Females who lost their mother early in life, but were high social status in adulthood (1SD above the mean) had a 5% lower hazard ratio compared to females who lost their mother but were of average social status. In contrast, females who lost their mother early in life, but were low social status in adulthood (1SD below the mean) had 20% higher hazard ratios, compared to the effects of maternal loss for females with average social status.

## Discussion

Previous work has debated the relative importance of early adversity and adult social relationships in determining survival in humans (*13, 20-22*). Our results shed light on this debate by providing an example of a wild animal model in which both early life experiences and adult social relationships are important and act independently on survival, with effects of similar magnitude. In addition, our moderation analysis indicates that at least for some sources of adversity, social relationships in adulthood may act as sources of resilience, allowing individuals to buffer the negative effects of poor early life experiences. Below, we consider several implications of these results, including the puzzle of weak mediation in spite of significant links between treatment, putative mediator, and outcome.

### The puzzle of weak mediation

We observed strong effects of both early adversity and adult behavior on survival, and effects of early adversity on at least one aspect of the adult social environment, with little or no mediation. One potential explanation for this set of observations is that an assumption of the mediation analysis was violated, thus producing spurious results. The most likely violated assumption is that of sequential unconfoundedness: i.e., if an unmeasured confounder in our system affects both the mediator and survival (*47, 48*). For example, individuals with better phenotypic or somatic quality (resulting from either genetic or environmental differences that were not included in our analysis) may experience both stronger social bonds and better survival, independent of early adversity. In this case, phenotypic/somatic quality would be an unmeasured confounder (see discussion of sequential unconfoundedness in *49*). To examine the potential for a confounding variable to affect our analyses, we conducted sensitivity analyses that assess how the mediated effect estimates vary as a function of the extent of the correlation between an unmeasured confounding variable and the mediator, and between that same variable and survival. Our sensitivity analyses demonstrate that our results are relatively robust to the assumption of sequential unconfoundedness (see Supplementary Text: “Sensitivity analysis for sequential unconfoundedness”; Figures S5, S6). As a consequence, it is likely that we are correctly estimating a small mediation size in this study.

A second, more likely, explanation is that the effects of early life adversity on social bonds in adulthood, albeit detectable, are relatively weak. Rosenbaum, Zeng, Campos, Gesquiere, Altmann, Alberts, Li and Archie (*34*) found similar results when testing for the mediating effect of social bonds for the relationship between early adversity and glucocorticoid levels. If early adversity does not have strong effects on social bond strength, then social bonds are unlikely to strongly mediate the comparatively quite strong connection between early adversity and adult survival.

In addition, our causal mediation pathway may be shaped by time-varying effects, as suggested by our simulation model. For example, if early life adversity affects social bonds early in adulthood, and survival is most strongly affected by social bond strength early in adulthood, then the matched timing of these effects could give rise to a strong mediating effect of social bonds. However, if early life adversity affects social bond strength earlier in adulthood, while survival is most strongly affected by social bond strength later in adulthood, then the mismatched timing of these effects would minimize the mediation effect. Previous work in birds and humans has shown that such time-varying effects may be a general phenomenon that warrants more attention (*7, 50, 51*). For example, in a survey of American adults, Nurius, Fleming and Brindle (*7*) show that social relationships in young adulthood are not linked to health, but that older adults with stronger social connections are in better health. Yang, Boen, Gerken, Li, Schorpp and Harris (*51*) also identified variability in the effects of social integration on several health biomarkers between American adolescents and adults. Exploring time varying effects of early adversity is therefore an important future avenue of exploration.

Two additional explanations are consistent with our observation of independent effects of social bonds and early adversity, combined with weak mediation. First, social bonds may be one of a larger set of mediators that all weakly mediate the link between early life environments and survival. Second, an as-yet unidentified variable could act as a strong mediator of early life adversity without involving social bonds. For example, the biological embedding hypothesis predicts that glucocorticoids – produced by the hypothalamic–pituitary–adrenal (HPA) axis and involved in regulating multiple physiological processes – link early adversity and lifespan (*52, 53*). In our study population, early life adversity predicts elevated concentrations of glucocorticoid metabolites in fecal samples in adulthood (*34*). Furthermore, elevated fecal glucocorticoid (fGC) concentrations in adulthood are associated with a shortened lifespan (*54*). At the same time, social bonds in adulthood are only modestly correlated with fGC concentrations (*34*), pointing to fGCs as a possible mediator of early life adversity that bypasses the pathway through social bonds. Notably, fGC concentrations, like social bonds, appear to weakly mediate the effects of early life adversity on survival (*40*), indicating that this pathway not only represents an alternative to the mediating pathway through social bonds, but also that multiple mediators may be involved.

### The evolutionary significance of sources of variance in survival

The independent effects of cumulative early life adversity and social bonds on female baboon survival are considerable. For each additional source of early adversity, lifespan is decreased by approximately 1.4 years, independent of social bond strength. Similarly, a one standard deviation decrease in social bond strength with either sex predicts 2.2 years of decreased lifespan, independent of early adversity. Notably, lifespan explains >80% of the variation in lifetime reproductive success (*26, 33, 55*), and females who experience early life adversity do not accelerate reproduction to compensate for the reduction in lifespan (*33*). Consequently, the combined effects of cumulative early life adversity and adult social isolation on survival have major consequences for lifetime reproductive success for female baboons.

These large effects on fitness indicate that phenotypes that allow individuals to survive in the face of multiple sources of adversity—which include features of the physical, social, and maternal environment—are likely to be favored by natural selection (*56-58*). Features of the social and maternal environment can be under direct natural selection. For example, our results suggest that selection should favor low adult mortality in part because maternal mortality directly decreases offspring survival in adulthood (in addition to other effects, such as the increase in the number of reproductive opportunities that comes with longer lifespans). In contrast, features of the physical environment (e.g., drought) cannot be under direct natural selection. However, adverse physical environments impose natural selection that acts on individual responses to environmental adversity. Indeed, work in humans has identified many genetic variants that influence the response to environmental stressors (e.g., pathogens, chemical stimuli), and some of these variants also carry genetic signatures of selection (*59, 60*). Thus, we expect natural selection to favor phenotypes that confer resilience to early life adversity even if the resulting phenotypes have lower fitness than phenotypes produced under advantageous early life conditions (*56-58*).

Adult social relationships also had strong and independent effects on adult survival, indicating that adult social behavior is not merely a proxy for the early life environment but is likely directly targeted by natural selection. Previous work on the links between social bonds and fitness did not control for early life experience (*35, 36*), limiting the ability to disentangle direct and indirect effects of adult behavior on fitness (*9, 61*). Our results suggest that adult social behaviors that maintain social bonds should be under strong selection. Further, because social behavior is almost always partially heritable (e.g., (*62-64*), these behaviors have the potential to evolve via natural selection. Further, they suggest that indirect genetic effects, in which the genotypes of social partners affect behavior, could play an important role in social selection and evolution (*65, 66*). Indirect genetic effects are unique because they illustrate that the environment itself can evolve and as a result create feedback loops that amplify or constrain evolutionary change, even in the absence of direct selection. However, selection on sociality is also likely to be limited by tradeoffs (*67, 68*). For example, tradeoffs may occur between the time allocated to sociality versus to other activities that are important for maintenance, such as foraging. In addition, sociality itself imposes costs, including potential increases in pathogen transmission, intraspecific competition, and social stress. Finally, the mechanisms that link adult social relationships to survival remain unclear, making it difficult to definitively identify potentially important targets of selection in addition to social bonds themselves.

### Individual sources of early adversity

We found strong effects of two individual sources of adversity on adult social bond strength and survival: maternal loss and drought. Consistent with previous findings (*26, 33, 34, 40, 69*), females whose mothers died when they were young had weaker social bonds with other females and reduced survival compared to females who did not experience early maternal loss, although they exhibited no differences in social relationships with males. In nonhuman primates, maternal loss during the juvenile period compromises the learning of social and foraging skills (*70-73*). In our study system in particular, losing a mother early in life is associated with shorter adult lifespans, weaker adult social bonds with females, compromised patterns of adult rank acquisition (*74*), elevated glucocorticoid concentrations in adulthood (*34, 39, 40*), and relatively poor survival of offspring (*69, 75*). Maternal loss also has negative consequences for adult phenotypes and fitness in other mammal species (*27, 75-79*) including humans (*80, 81*). Therefore, maternal loss during development represents a strong source of early adversity across taxa, especially in species where mothers are essential for the development of crucial skills.

In addition to maternal loss, drought emerges as an important source of early life adversity in this analysis. Females who experienced drought in their first year of life had weaker social bonds with both females and males, and also experienced reduced survival relative to females born in non-drought years via both mediated and direct effects (Tables 2, 3). Drought threatens food availability which in turn hinders growth and development during the crucial first year of life (*82-86*). In addition, individuals born during drought may have fewer opportunities to learn foraging skills during younger years when adults are more tolerant of them during foraging (*72, 87*). Consistent with our results, experiencing dry seasons and droughts in early life negatively affects health in humans (*88-93*).

Notably, two previous analyses in our study system found that drought did not predict adult survival independently of other sources of early adversity (*26, 33*). The difference between the previous studies and this one may be attributable to using somewhat different subsets of the long-term data, because of different data requirements for each analysis. For instance, the current analysis includes a larger representation of females who were born during a particularly severe drought in 2008-2009, a two-year consecutive period in which annual rainfall was less than 200 mm (*94*). This drought inflicted substantial mortality on wildlife and livestock throughout the Amboseli ecosystem and surrounding areas (*95, 96*). Therefore, it represented an extreme climatic event in the early lives of these individuals which may have driven the strong effects of drought not detected in previous analyses (*26, 33*).

### Moderating effects of adult behavior

Our analyses indicate that strong social bonds in adulthood may buffer the negative consequences of adverse early life events—even for maternal loss, which has far-reaching effects on phenotypes and fitness. Specifically, we found that female baboons who lost their mothers in early life but were able to maintain strong social bonds in adulthood survived better than those who lost their mother and were socially isolated. Therefore, resilient adult phenotypes may buffer the negative effects of maternal loss. Social buffering has also been suggested as a mechanism to counteract the negative effects of early life adversity in other mammals (*18, 97*) and humans (*81*). For example, mountain gorillas who lose their mothers tend to strengthen their social bonds with other group members; perhaps as a consequence, they suffer no detectable survival costs from maternal loss (*97*). Social bonds with males may be a particularly important buffer as, unlike social bonds with females, they are not weakened by maternal loss (*34*).

### Conclusions and future directions

By linking prospective data on early life adversity with data on social bonds and survival in adulthood, we find support for both social causation and health selection. Specifically, by accounting for the complex relationships between early life, adulthood, and survival, we confirmed the far-reaching effects of early life adversity – which contributes directly to both compromised adult social relationships and adult survival – and we also confirmed a direct influence of adult social relationships on survival. Furthermore, for at least some sources of early adversity, strong adult social bonds can reduce the negative effects of early life adversity. In addition to finding support for both social causation and health selection, we argue that responses to early adversity, sources of early adversity, and adult social behavior are all likely targets of natural selection. Future work should explore how variation in the timing of early life effects, and in the timing of the effects of adult phenotypes, affect connections between early adversity, mediators, and survival in other species. Future work should also examine other potential mediators (e.g., phenotypic quality, immune response, glucocorticoid levels) of the relationship between early adversity and lifespan.

## Materials and Methods

### Study Subjects

We used longitudinal data on 199 wild adult female baboons (*Papio cynocephalus*, with some natural admixture from the closely related species *P. anubis* (*98, 99*) from the Amboseli ecosystem in Kenya collected between 1983 and 2019. Subjects are habituated to and individually recognized by experienced observers who collect demographic and behavioral data 6 days a week, year-round, following 1-2 social groups (‘study groups’) per day. Birth and death dates for all study subjects are accurate to within a few days’ error. Two original study groups (studied beginning in 1971 and 1980 respectively) experienced multiple permanent fissions and fusions over the years, resulting in a total of 19 different social groups that persisted for varying lengths of time. Female baboons remain in their natal social group throughout their lives (except for group fissions or fusions), and thus any disappearance of a female in our dataset was considered a death. Of the 199 females in the study, 74 had died by the end of the study and the rest were considered censored in survival analyses. To be included in the study, females had to meet the following criteria: (i) they survived to at least 4 years of age (most females reach menarche between 4 and 5 years of age; (*100*), (ii) they had available data on exposure to all six sources of early adversity in the infant and juvenile period, and (iii) they were members of study groups that foraged entirely on naturally occurring foods (*26, 33, 34*).

### Measuring Early Life Adversity

We created an index of cumulative early life adversity by considering six conditions that represent socioenvironmental adversity experienced during the first four years of life: drought in the first year of life, large group size at birth, low maternal social status at birth, low maternal social connectedness in the first two years of life, a close-in-age younger sibling, and maternal loss before age four (Table 1; *26, 33*). Drought years were those in which less than 200 mm of rain fell. Large group sizes were considered as those in the highest quartile of the group size (number of adults) distribution. Low maternal social status was assigned when the mother’s proportional dominance rank in the month of her offspring’s birth was in the lowest quartile of dominance ranks. Proportional dominance rank ranges from 0 (lowest ranking female) to 1 (highest ranking female) and indicates the proportion of adult females in a study subject’s social group that she dominated in agonistic interactions (*46*). Low maternal social connectedness was assigned when the mother’s social connectedness to other females was in the lowest quartile of the distribution of social connectedness values during our study subjects’ first two years of life. Following previous work on early life adversity in this population (*26, 33*), social connectedness was measured as the relative frequency of the mother’s grooming interactions with other adult females in her social group, adjusted for observer effort (see ‘Measuring social bond strength’ for information about observer effort). Close-in-age younger siblings were those born within 1.5 years of the subject’s birth, approximately the shortest quartile of observed interbirth intervals in the Amboseli baboons (*26*). A subject was considered to experience maternal loss if her mother died within her first four years of life (i.e., before the earliest age of sexual maturation for females in this population).

Each subject’s cumulative adversity index was calculated as the sum of exposures to these six sources of adversity. In our dataset, 48 females experienced zero sources of adversity, 84 experienced one, 50 experienced two, 14 experienced three, 3 experienced four, and none experienced five or six.

### Measuring Social Bond Strength

We measured an adult female’s social relationships by assessing the strength of social bonds with her top three male or female social partners separately, in each year of her life, measured relative to the social bonds of all other females in the population with males or females respectively, as described in Rosenbaum, Zeng, Campos, Gesquiere, Altmann, Alberts, Li and Archie (*34*). Briefly, grooming interactions are recorded during all hours of observation, using representative interaction sampling in which observers record all the interactions they see while conducting 10-minute focal follows on a randomized set of individuals. We calculated the number of grooming interactions with each partner per day of co-residence in the same group from these representative interaction data for each year of life for each female subject starting on her birthday. Calculating interaction rates from such data is complicated by the fact that the number of observers remains constant over time, while social group sizes vary, so that higher numbers of grooming interactions per pair of animals (per dyad) will generally be observed in smaller groups compared to larger groups. We corrected for this variation in observer effort by regressing daily rates of grooming interactions per dyad against observer effort, where observer effort was calculated as the number of focal samples on adult females collected during each observer day, divided by the mean number of adult females in the group during those days, divided by the number of days that each dyad was co-resident (*34, 36*). We z-scored the corrected rates within years to control for temporal variation in sociality in the population.

Each subject’s social bond strength with females and with males was taken as the average of the subject’s three strongest adult female grooming partners and adult male grooming partners, respectively, to calculate a dyadic sociality index (DSI). A positive value for social bond strength indicates a female had relatively strong social bonds with her top three partners compared to the population average. A negative value for social isolation means the subject had relatively weak social bonds with her top three partners.

### Random Effects and Covariates

Previous work has demonstrated that several environmental and demographic variables not discussed above (i.e., presence of maternal relatives, group size, social status, percent of prior year with young infant, percent of prior year cycling, rainfall) explain variation in social bond strength and/or survival (*34-36, 42*). To control for these effects, we included them as covariates in our mediation and moderation analyses (for details see Supplementary Information). We also included social group and hydrological year as random effects in all models to control for group-to-group and interannual variation (*34*). Age was not included as a covariate even though social bonds vary with age, because age effects are captured by our functional principal components analysis (FPCA) approach to modeling the mediator (see below). Because our baboon study population represents an admixed population (yellow baboon ancestry is dominant, but all individuals exhibit some degree of admixture with anubis baboons), we also ran separate analyses that included a covariate measure of individual admixture, a ‘genetic hybrid score’ that represents the proportion of each individual’s genome estimated to be from *P. anubis* ancestry (see Supplementary Information, also (*101, 102*). Results that incorporated hybrid score (Tables S4-S5) were similar to those of the full model (Tables 2-3).

In preliminary analyses we considered social status as a third potential mediator of the effects of early adversity on survival. However, as previously reported (*35, 36*), we found no effects of social status (again measured as proportional dominance rank) on female survival (Table S6). In addition, we found no effect of cumulative early adversity on female social status, and no mediating effects of female social status on the relationship between early life adversity and survival (Tables S6). As a consequence, we focus on social bond strength as the primary mediating variable in the main text, but report models for social status as a mediator in the Supplementary Materials.

One individual source of early adversity strongly predicted proportional dominance rank: low maternal dominance rank predicted low proportional rank for the study subject in adulthood (Table S6), which is unsurprising as rank is matrilineally inherited in this species (*103*). In light of this relationship, we controlled for proportional rank by including it as a covariate when estimating the effect of early life adversity on the mediator.

### Mediation Analysis Implementation

We fit two models in each of our mediation analyses (*40*). The first model captures the relationship between early adversity and the mediator. The second model characterizes the relationship between early adversity, the mediator, and survival. Models were implemented using the R packages survival and flexsurv. The reproducible code is available at [https://github.com/zengshx777/MFPCA_Codebase].

#### The first model: the relationship between early adversity and the mediator

Our first model applies to the observed mediator trajectory *M*_*ij*_ and the measure of early adversity *A*_*i*_, where *i* indexes individual and *j* indexes time. This model corresponds to the Equation 1 in the main text. Because the observed mediator values are noisy and potentially measured imprecisely, we consider them, after adjusting for covariates and random effects, as realizations of an underlying smooth process (M_i_(*t*_*ij*_)) with a random noise. Specifically, we modeled the trajectory of the mediator *M*_*ij*_ as a combination of covariate effects *C*_*ij*_*β*_*m*_, social group random effects 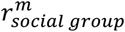, hydrological year random effects 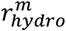, an underlying smooth process M_i_(*t*_*ij*_), and an error term *ε*_*ij*_,

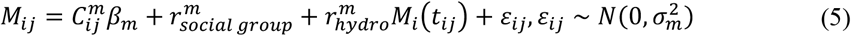

Because M_i_(*t*_*ij*_) is of infinite dimension mathematically, we performed dimension reduction to improve the statistical power of our analysis. Specifically, we used a functional principal component analysis (FPCA) method to decompose the smooth process as the linear combination of the fewest possible functional principal components (*39, 40, 104-106*). We began by examining the correlation between any two time points in the mediator process (e.g., between the value of the mediator at age 4 and age 8, between the value of the mediator at age 4 and age 9, and so on) to produce a correlation structure between mediator values at different time points, which we then expressed as principal components or eigen functions,

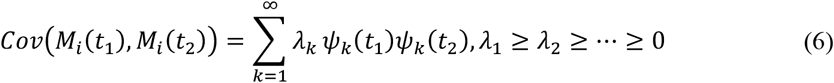

where *λ*_*k*_ is the explained variance of the orthogonal normal principal components *Ψ*_*k*_(*t*). We ordered the principal components by the amount of variance they explained to reflect the fact that principal components that explain more variance (larger *λ*_*k*_) are more important in expressing the smooth process. We then used the first *K* principal components, where *K* is the number of components necessary to collectively explain at least 90% of the variance 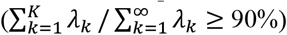.

In the next step, we represent the smooth process of each subject’s mediator process as a linear combination of the *K* principal components,

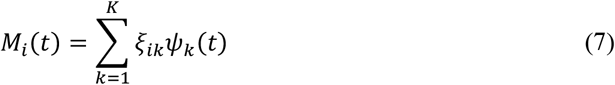

where *ξ*_*ik*_ is the principal score for individual *i* on the *k*th principal component or eigen function. The variance of *ξ*_*ik*_ corresponds to the explained variance of principal component, *λ*_*k*_. We can efficiently express the smooth process and trajectory with a small number of principal components (*K* is never greater than 4 in our work), capturing the major variation. Therefore, coupled with the FPCA, we posit the following model of the mediator,

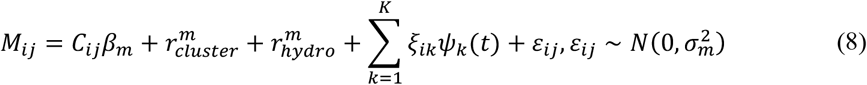

which corresponds to Equation 1 in the main text. Furthermore, we assume that the differences in trajectories caused by early adversity are captured by the differences in the principal scores. Therefore, we use the following specification for the principal scores, with different means for each level of adversity in the cumulative model or with different means for the group that experienced each early adversity and for the group did not experience early adversity in the models of individual sources of adversity,

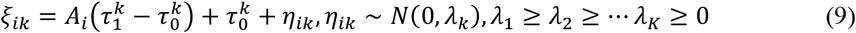

Where 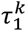 denotes the mean of the k^th^ principal score for the subjects in the adversity group while 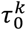 represents that for the non-adversity group (for cumulative adversity, 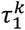 denotes the k^th^ principal score for a higher level of adversity relative to 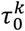, the score for one level lower) We fit Equation 8 simultaneously with Equation 9. Hence, instead of estimating the effect of adversity on the trajectories directly, which is a high dimensional problem, we estimate its effect on the first K principal scores *ξ*_*i*1,_ *ξ*_*i*2,_ ⋯, *ξ*_*iK*_.

The effect of early adversity parameterized with different means for the principal scores is not directly interpretable. Therefore, we estimated the effect of early adversity on the mediator as the difference in the mean of the trajectories for the adversity group versus the non-adversity groups (for the cumulative adversity measure it was the difference in means comparing two adjacent levels of adversity, e.g., for a cumulative score of 3 versus 2). Based on Equations 8 and 9, we can express the conditional expectation of the mediator process *M*_*ij*_ at time point *t*_*ij*_ as follows,

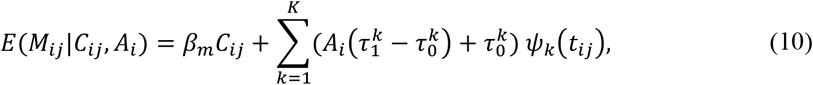

which corresponds to Equation 1 in the main text. Next, we express the effect of early adversity on social isolation using:

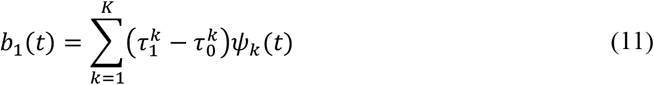

The effect on the mediator is also time-indexed, because we are estimating the effect of adversity on the mediator trajectory across the lifespan. Integrating *b*_1_(*t*) over time gives an estimation of parameter *β*_1_(the beta coefficient associated with the effect on the mediator) in Equation 1 in the main text:

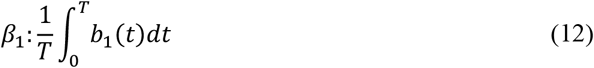

#### The second model: the relationship between early adversity, the mediator, and survival

Our second model estimated the survival outcomes. We adopted a Cox model for the hazard rate *λ*(*t*). Specifically, we employed the following model,

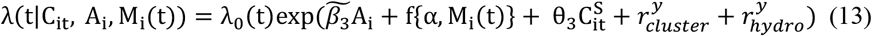

where (i) f{α, M_i_(t)} is the function of the mediator process up to time point *t* with parameter *α* characterizing the effect of the mediator process on the hazard rate, and M_i_(t) is replaced by its estimated value 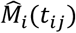 from Equation 8, and (ii) *λ*_0_(t) is the baseline hazard rate, which we specify as following a Gompertz distribution (*107, 108*),

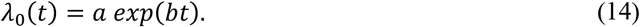

We consider two specifications of *f* in our case: (i) a model using estimated trajectories of three-year mediator values that assumes the hazard rate depends on the mediator history in the previous three years, 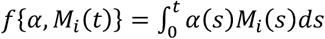, and (ii) a model using estimated trajectories of one-year mediator values that assumes the hazard rate depends on the current mediator value assessed in the year in which survival is assessed, f{α, M_i_(t)} = *α M*_*i*_(*t*). For the three-year model, we specify *α*(*t*) as a linear combination of the spline basis *α*(*t*) = *s*(*t*)′*ρ, s*(*t*) = [1, *t*, (*t* − *k*_1_) ^2^, (*t* − *k*_2_) ^2^,.., (*t* − *k*_*L*_) ^2^] (*106*), which allows a flexible modeling of how the past mediator affects the survival.

Following the notation in the causal mediation analysis literature (*109, 110*), let *SZ, Z*^′^(*t*) denote the survival function when the subject’s early adversity status is *Z* and the mediator trajectory counterfactually takes the value as if the subject has early adversity status *Z*^′^. The adversity status *Z* can be ordinal (for cumulative adversities) or binary (*z*=0 for the non-adversity group and *z*=1 for the adversity group mediator to estimate the total, direct, and mediated effects). For example, if *z*=0 and *Z*^′^=1, then *S*_*Z,Z*_^′^(*t*) is the survival function for baboons who did not experience early adversity, but whose mediator values are counterfactually calculated as if they did experience early adversity. This strategy is standard in causal mediation research; it allows us to decompose the total effect into the mediated effect and the indirect effect (*40, 110-112*). Based on the model for hazard rate, we can calculate the *S*_*Z,Z*_^′^(*t*) up to time *t* by integrating the hazard function. Specifically, it takes the following form,

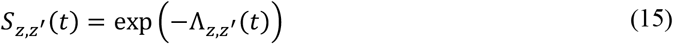

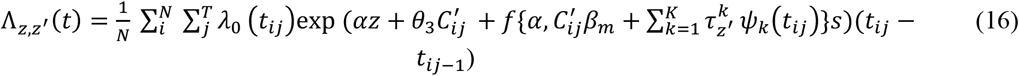

where Λ_*Z,Z*_^′^(*t*) is the cumulative hazard function. Once we obtain *S*_*Z,Z*_^′^(*t*), we can calculate the total effect, direct effect, and mediated effect on the scale of years,

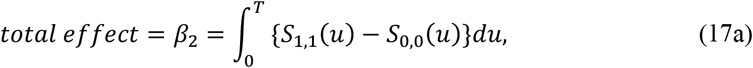

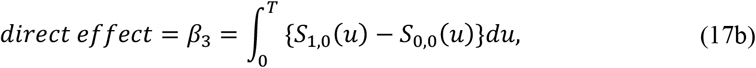

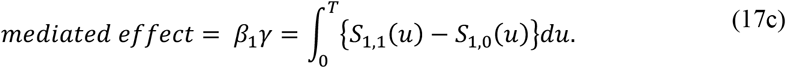

To estimate the effect of the mediator on survival (while controlling for the effects of early adversity on the mediator), we followed similar steps. We calculated the mediator trajectory for a one unit change in social bond strength while fixing the value of early adversity exposure to one (a value that corresponds to experiencing at least one source of adversity in the cumulative adversity model or to experiencing adversity in the models for each individual source of adversity). This approach allows us to estimate the consequences of the unit change in the mediator, irrespective of the underlying reason why it might change (i.e., whether due to the effects of early adversity or some other reason), because it controls for the effects of early adversity that act independently of the mediator. Thus, the isolation effect describes how one unit change in social bond strength affects survival in years, where a one unit change represents approximately 1.7 SD for social bond strength with females and 1.4 SD for social bond strength with males; 1 SD in social bond strength with females=0.59 social bond strength units, 1 SD in social bond strength with males=0.70 units.

#### Causal Assumptions

To interpret the above models as causal, three assumptions are required. The first is the assumption of unconfoundedness. In our case, we assume that early adversity is randomly assigned to the subjects in the study. It also assumes that no unmeasured confounding variables cause variation in both early adversity and the mediator or cause variation in both early adversity and survival time, a result that follows if exposure to early adversity is largely determined by natural events that are independent of the subject’s individual traits, which is most likely true in our case.

The second is the assumption of sequential unconfoundedness, which states that no unmeasured confounding variables cause variation in both the mediator and survival, besides the observed covariates *C* and the past history of the mediator *M* (*47, 48, 113*). This assumption will be violated if an unmeasured variable (for instance, phenotypic or somatic quality, resulting from either genetic or environmental differences that were not included in our analysis) enhances or reduces both the mediator and survival.

We controlled for confounders as much as possible by including covariates when modeling the mediators and survival, but the sequential unconfoundedness assumption is essentially untestable because it invokes the possibility of an unknown and therefore unidentified covariate (*49*). To estimate the potential effect of one or more unidentified covariates, we performed a sensitivity analysis (for details see Supplementary Text: “Sensitivity analysis for sequential unconfoundedness”). Specifically, we assumed the existence of an unmeasured confounder between the mediator and survival that violates the sequential unconfoundedness assumption (*114, 115*). In our simulation, the correlation between the unmeasured confounder and the mediator or outcome quantifies the degree of violation of the assumption. For a set of prespecified correlation values, we repeated the mediation analysis and examined the sensitivity of the results to the degree of violation of the sequential unconfoundedness assumption. We found that under various degrees of violation of the assumption, the mediated effect was not significant (Figures S5, S6). Therefore, our conclusions are robust to the untestable assumption. Details of the sensitivity analysis can be found in the supplement and Zeng, Lange, Archie, Campos, Alberts and Li (*40*).

The third assumption we impose is independent censoring, i.e., we assume that the time at which a subject drops out of the study prior to death is random with respect to characteristics of the subject or its experience of early adversity. This assumption is likely to hold in our study because female baboons are censored in only two circumstances in our study: either they survived to the end of the period of data collection, or the social group in which they lived was dropped for logistical reasons.

### Moderation Analysis Implementation

For the moderation analysis, we modified Equation 13 by incorporating an interaction term between *A* and *M* in the hazard function for survival, as follows:

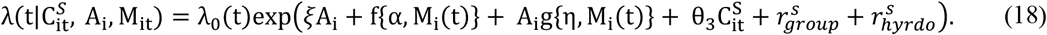

Adding this interaction term, A_i_g{η, M_i_(t)}, in the hazard function allows us to test for the interaction between early adversity and social behavior predicted by the social buffering hypothesis. Therefore, this approach allows us to estimate how the effects of early adversity on survival vary across different levels of the moderator. Similar to the survival model in the mediation analysis, we imposed two specifications for the interaction term A_i_g{η, M_i_(t)}: (i) a three-year model 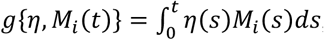, and (ii) a one-year model, g{η, M_i_ (t)} = *η M*_it_. For the three-year model, we use 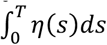 as the summary for the moderation effect. When *η* < 0, the model indicates that a higher value for the moderator buffers the negative effects of early adversity. When *η* > 0, the model implies that the moderator amplifies these negative effects.

## Acknowledgments

In Kenya, our research was approved by the Kenya Wildlife Service (KWS), the Wildlife Research & Training Institute, the National Environment Management Authority (NEMA), and the National Council for Science, Technology, and Innovation (NACOSTI). We also thank the University of Nairobi, the Institute of Primate Research, the National Museums of Kenya, the members of the Amboseli-Longido pastoralist communities, the Enduimet Wildlife Management Area, Ker & Downey Safaris, Air Kenya, and Safarilink for their cooperation and assistance in the field. Particular thanks go to the Amboseli Baboon Research Project field team (R.S. Mututua, S. Sayialel, J.K. Warutere, I.L. Siodi, G. Marinka, B. Oyath) and camp staff. We also thank T. Wango and V. Oudu for their untiring assistance in Nairobi, and Jeanne Altmann for her fundamental contributions to the Amboseli baboon research. The baboon project database, BABASE, was designed and programmed by K. Pinc and is expertly managed by N.H. Learn and J.B. Gordon. For a complete set of acknowledgments of funding sources, logistical assistance, and data collection and management, please visit http://amboselibaboons.nd.edu/acknowledgements/.

## Author contributions

Conceptualization: ECL, FL, EAA, SCA

Data Curation: JT, EAA, SCA

Formal analysis: SZ, FL

Funding Acquisition: FL, JT, EAA, SCA

Methodology: ECL, SZ, FAC, FL, EAA, SCA

Visualization: ECL, SZ

Resources FL, JT, EAA, SCA

Supervision: FL, SCA

Writing—original draft: ECL, SZ

Writing—review & editing: ECL, SZ, FAC, FL, JT, EAA, SCA

## Data and materials availability

The reproducible code is available at [https://github.com/zengshx777/MFPCA_Codebase]. All data used in this study will be archived on Duke University Digital Repository.

