## Supplemental Materials for "Early life adversity and adult social relationships have independent effects on survival in a wild animal model of aging"

### Supplementary Text

<u>Results that recapitulate previous analyses</u>

*Total and direct effects ($\beta_2$ and $\beta_3$) of individual sources of adversity: maternal loss and drought reduce survival*

Among the six individual sources of early adversity, maternal loss and drought exerted the strongest and most consistent effects on adult female survival (Tables 2-3; Figures S1-S2; these results are similar but not identical to the results reported for early adversity by *26*). Females that lost their mother before four years of age had adult lifespans that were reduced by 3.3 years (total effect, $\beta_2$, black brackets in Figure 1C). In the current analysis, more than 80% of the effect of maternal loss on survival can be explained by effects outside the pathways that include adult social bonds (direct effect, $\beta_3$, green arrows in Figure 1C; the estimate varied from 2.67 to 3.21 years across the two mediation models). Females that experienced drought in the first year of life lost 2.70 years of life relative to those that did not experience drought (again, the estimate varied across the two models), and like other measures of adversity, most of the effect of early life drought on survival was direct and outside the pathway that includes adult social bond strength (direct effect, $\beta_3$, green arrows in Figure 1C).

*Effects on the mediators ($\beta_1$): Early life adversity weakens social bonds with females, but not social bonds with males*

Cumulative early adversity, as well as most individual sources of early adversity, resulted in weaker social bonds with adult females (Table 2, orange arrow in Figure 2A; Figure S1; recapitulating cumulative adversity results from *26*). The effect size for cumulative early adversity was relatively weak: each additional source of cumulative adversity reduced social bond strength with females by 0.09 “social bond strength units,” (see Materials and Methods) which is equal to 15% of 1 SD in social bond strength with females (1 SD in social bond strength with females=0.59 social bond strength units). Among the individual sources of adversity, drought, close-in-age sibling, maternal loss, and low maternal social status were all associated with weaker social bonds with adult females (Table 2, Figure S3). As with the direct effects of early adversity on survival, early life drought and maternal loss had the strongest effects on social bonds with females; drought was associated with an average decrease in bond strength with adult females of 0.21 social bond strength units (36% of 1 SD), while maternal loss decreased bond strength with adult females by 0.26 units (44% of 1 SD).

Cumulative adversity did not predict weaker social bonds with adult males (recapitulating results from *26*), although two individual sources of adversity – early life drought and low maternal social status – did (Table 3; orange arrow in Figure 2B; Figure S2). Females with low-ranking mothers experienced a decrease in social bond strength with adult males of 0.25 units (36% of 1 SD; 1 SD in social bond strength with males=0.70 units), while those that experienced early life drought showed a decrease in social bond strength with adult males of 0.16 units (23% of 1 SD). In contrast to the strong effects of maternal loss on social bond strength with adult females, maternal loss did not predict social bond strength with adult males: the estimated effect size was near zero (Table 3; Figure S3).

Covariates for mediators and survival

Based on previous analyses, we controlled for known environmental effects on the mediators and on survival. As described in Rosenbaum, Zeng, Campos, Gesquiere, Altmann, Alberts, Li and Archie (*34*), when modeling variation in social bond strength with females and males, we included mean group size (number of adults, mean 27.6 ± 0.21 s.e. individuals; range 5-45 individuals), mean number of co-resident adult maternal relatives (mothers and sisters; mean 1.68 ± 0.03 s.e.; range 0-7), and mean proportional dominance rank (mean 0.51 ± 0.007 s.e.; range 0-1) as model covariates. We also included proportion of prior year with young infant (<3 months of age) for social bond strength with females (mean 0.11 ± 0.003 s.e.; range 0-0.83) and proportion of prior year cycling for social bond strength with males (mean 0.35 ± 0.007 s.e.; range 0-1). All covariates for the social bond models were measured on a yearly basis.

In the survival models, we included group size, group size squared, proportional dominance rank, number of co-resident adult maternal relatives, and annual delta rainfall (mean -11.3 ± 3.24 s.e. mm; range -267-415 mm) as covariates. Annual delta rainfall is calculated by averaging the rainfall during a given year window across all years with data and subtracting this average from the mean rainfall from the year of life window (*34*). All covariates in the survival model were measured on a yearly basis and those that overlap with social bond covariates have the same mean and range (see above). We also included random effects of social group and hydrological year in all models.

In a subset of analyses, we also included an effect of hybrid score on the mediators and on survival to assess the possible contributions of genetic admixture. Hybrid score measures the proportion of each subject's genome estimated to be from *P. anubis* ancestry based on microsatellite markers, such that a score of 1 corresponds to unadmixed *P. anubis* ancestry and a score of 0 corresponds to unadmixed *P. cynocephalus* ancestry (*101, 102*). Using hybrid score as a covariate reduced our sample size to N=165 females because not all subjects have genetic information. Therefore, we report results from models using hybrid score as a covariate in the supplemental section *Effects of hybrid score* below (Tables S4-S5).

Data imputation strategies for missing data

Data were missing due to our exclusion of data collected during low observation periods and group fission events. Based on preliminary analyses, to recover 80% accuracy relative to a full year of data, we needed at least 9 months of time with infant data, 7 months of cycling and social bond data, and one month of rank, group size, and maternal kin data. Covariates from subjects who were observed for less than these 80% cutoffs were considered missing.

One or more covariates were missing from the dataset used to model social bond strength with females in 121 out of 1849 years of life and from social bond strength with males in 140 out of 1849 years of life. Females were also missing social bond strength values in 229 out of 1849 years of life for social bond strength with females, and 253 out of 1849 years of life for social bond strength with males. The dataset used for survival contained missing values for one or more covariates in 58 out of the 1849 female years of life.

Missing covariates or social bond strength values for a subject's year of life were imputed based on two strategies. For group size, the number of adult maternal kin, and proportional rank in the social bond strength and survival datasets, we used the average of the previous year and subsequent year's values to replace missing data; this strategy worked well for group size, female group membership, and female proportional rank because they are relatively stable over time. For all other covariates and social bond strength values, we employed a random age-specific imputation procedure as in Campos, Villavicencio, Archie, Colchero and Alberts (*36*).

Simulation to explore the weak mediated effect

Our estimated mediated effects in the mediation analyses were small despite significant effects of early adversity on the mediators and the mediators on survival. We hypothesized that the small mediated effect resulted from differences in timing of effects (e.g., that the effects of early adversity on the mediator differ temporally from the effects of the mediator on survival). To test the plausibility of this hypothesis, we designed a simulation with two stages ($T$ = 1,2) which represent two different periods in the life course (e.g., early and late adulthood). To capture the effects of early adversity on social bond strength observed in the baboon system, we assigned values of adversity ($A_i$), randomly across individuals as $A_i \sim Bern(0.5)$); we specified that higher values of adversity were associated with lower values of the mediator ($M_i$), and we specified that higher values of the mediator were associated with higher chance of survival. We then varied whether the effects of adversity on the mediator occurred at stage 1 or stage 2, and whether the effects of the mediator on survival occurred at stage 1 or stage 2.

To capture time varying effects of early adversity on the mediator at the two different time stages, we modelled the mediator as:

$$\left.\begin{array}{l} M_{1i} \sim Bern(0.55 - 0.3A_i) \\ M_{2i} \sim Bern(0.75 - 0.1A_i) \end{array}\right\} if\ A_i\ has\ strongest\ effect\ on\ M_i\ early\ in\ adulthood\ (stage\ 1), \quad (S1a)$$

$$\left.\begin{array}{l} M_{1i} \sim Bern(0.55 - 0.1A_i) \\ M_{2i} \sim Bern(0.75 - 0.5A_i) \end{array}\right\} if\ A_i\ has\ strongest\ effect\ on\ M_i\ late\ in\ adulthood\ (stage\ 2). \quad (S2b)$$

Note the difference between the effect size of early adversity on the mediator in early versus late adulthood is larger in equation S1b than S1a because fewer individuals survive to late adulthood; this approach maintains a similar overall effect of early adversity on the mediator across life stages. Survival probability to stage $T$ only takes the values 0, 1, and 2. All subjects survived to stage $T$=0; $T=1$ indicates the subject died at stage 1; $T=2$ indicates the subject died at stage 2. Survival probability was equal to:

$$P(T \geq 1) = 0.8 - 0.1A_i + 0.2M_{1i}, for\ survival\ to\ at\ least\ stage\ 1, \quad (S2a)$$

$$P(T \geq 2) = 0.1 - 0.1A_i + M_{1i}\gamma_1 \mp M_{2i}\gamma_2, for\ survival\ to\ at\ least\ stage\ 2\ , \quad (Sb2)$$

Where $\gamma_1 \geq 0$, $\gamma_2 \geq 0$, and $\gamma_1 + \gamma_2 = 0.5$, such that $\gamma_1$ represents the strength of the effect of the mediator at stage 1 on survival to at least stage 2 and $\gamma_2$ represents the strength of the effect of the mediator at stage 2 on survival to at least stage 2. The effect of early adversity on the mediator, the mediator on survival, and the mediated effect were calculated using Monte Carlo methods. We fixed the effect of early adversity on the mediator and the effect of the mediator on survival in order to examine how altering the timing of these effects alone alters the strength of the mediated effect.

Our simulations support the idea that the strength of the mediated effects is determined by the timing of the effect of early adversity on the mediator combined with the mediator on survival (Figure S4). Note that in our simulations we fixed the effect of early adversity on the mediator, as indicated by the fixed values of $M_i$ in equations S1a and S1b ("Effect on mediator", orange arrow in Figure 1C) as well as the effect of the mediator on survival, as indicated by the fact that $\gamma_1 + \gamma_2 = 0.5$ ("Bond effect", purple arrow in Figure 1C). Even when these values were fixed, we found that the estimate of the mediated effect (pink arrows in Figure 1C) depends on the timing of these effects. The largest mediated effects were observed when the timing of these two effects is matched such that either (i) early adversity affects the mediator early in adulthood and survival depends most strongly on the mediator's value early in adulthood, or (ii) early adversity affects the mediator late in adulthood and survival depends on the mediator's value late in adulthood.

<u>Sensitivity analysis for sequential unconfoundedness</u>

We performed sensitivity analyses on the second of the three assumptions required to interpret causality in our models, the assumption of sequential unconfoundedness (see 'Causal Assumptions' in the Methods section). Sequential unconfoundedness assumes that no unmeasured confounders at one stage of life affect both the mediator and survival in the next stage of life (*47, 48, 113*). The sequential unconfoundedness assumption is crucial for identifying effects in the mediation analysis (e.g., the mediated effect, direct effect, and bond effect), yet it is generally untestable with observed data (*49*).

To test the sensitivity of our models to this assumption, we posited a hypothetical unmeasured confounder that would violate the sequential unconfoundedness assumption, and examined how our estimates varied depending on the degree of violation (*114, 115*). Specifically, we introduced a binary unmeasured confounder $U_i$ that was negatively correlated with both the mediator and the survival. We expanded the model for the mediator (Equation 8) and survival (Equation 13) as:

$$M_{ij} = C_{ij}\beta_m + r^{m}_{cluster} + r^{m}_{hydro} + \sum_{k=1}^{K} \xi_{ik}\psi_k(t) + \zeta_M U_i + \varepsilon_{ij}, \varepsilon_{ij} \sim N(0, \sigma_m^2) \quad (S3)$$

$$\lambda(t|C^{S}_{it}, A_i, M_{it}) =$$

$$\lambda_0(t)\exp(\beta_3 A_i + f\{\alpha, M_i(t)\} + g\{\eta, A_i M_i(t)\} + \theta_3 C^{S}_{it} + \zeta_S U_i + r^{s}_{group} + r^{s}_{hyrdo}) \quad (S4)$$

where $\zeta_M$ and $\zeta_S$ are the pre-specified sensitivity parameters that control the correlation between the unmeasured confounder and mediator and the survival outcome, respectively. When the sequential unconfoundedness assumption is valid, no unmeasured confounder is simultaneously correlated with the mediator and survival outcome, which implies $\zeta_M \zeta_S = 0$. When both $\zeta_M$ and $\zeta_S$ take non-zero values, the sequential unconfoundedness assumption is violated. Therefore, we can use $(\zeta_M, \zeta_S)$ as the sensitivity parameters to measure the degree to which the sequential unconfoundedness assumption is violated.

The sensitivity analysis takes the following steps. First, we specified a grid of values for the sensitivity parameters$(\zeta_M, \zeta_S)$. Specifically, we chose $(\zeta_M \in \{-0, -0.1, -0.2, -0.5, -1\}, \zeta_S \in \{0, 0.1, 0.5, 1\})$ to model potential unmeasured confounders in this system, because these values restrict the unmeasured confounder to be negatively correlated with the mediator and negatively correlated with survival. In the second step, with every fixed pair of $(\zeta_M, \zeta_S)$, we fit the mediator and survival models above. Compared with the original models (8) and (13), we needed to have an additional step of simulating the unmeasured confounder $U_i$ drawing from a Bernouli distribution from the observed data and $(\zeta_M, \zeta_S)$ and the other model parameters. Finally, we estimated the mediated effect with cumulative adversity based on the model with unmeasured confounders. We repeated the above steps with all possible combinations of $(\zeta_M, \zeta_S)$ on the pre-specified grid and examined how sensitive the estimates of the mediated effect are to the values of $(\zeta_M, \zeta_S)$, which reflects how sensitive the estimates are to the violation of sequential unconfoundedness. We consider our results to be robust to a violation to sequential unconfoundedness if the mediated effect in the sensitivity analysis is small even when the sensitivity parameters $(\zeta_M, \zeta_S)$ are varied. If those conditions are met, even if unmeasured confounders exist, they are unlikely to qualitatively influence our analysis or interpretation. Figures S5 and S6 summarize the results of the sensitivity analysis for the two mediators (social bond strength with females and males) under the pre-specified grid of $(\zeta_M, \zeta_S)$, with one-year mediator values and with three-year mediator values, respectively. We found that our mediated effect estimates were robust to violations to sequential confoundedness, making it unlikely that unmeasured confounders affect our results (Figures S5, S6). The point estimates of mediated effects generally become closer to zero as $\zeta_M$ decreases or $\zeta_S$ increases, and the size of the mediated effect becomes negligible when $\zeta_S \geq 0.5$ and $\zeta_M \leq -0.1$. In addition, the credible interval becomes wider when either one of the sensitivity parameters $(\zeta_M, \zeta_S)$ increases in magnitude. These patterns make sense as we specify the $U_i$ to be negatively correlated with the mediator and decrease the survival probability, which in turn reduces the proportion of the adverse effect that can be explained by the mediator.

When we used one-year social bond strength with either sex as a mediator, the mediated effects remain small and not significant across all possible magnitudes and combinations of correlations between the unmeasured confounder and the mediator $(\zeta_M)$ and between the unmeasured confounder and the survival outcome $(\zeta_S)$ , i.e. for all combinations of the sensitivity parameters ($\zeta_M, \zeta_S$; Figure S5). This sensitivity analysis further supports our results of weak mediated effects in the relationship between early adversity and survival, based on one-year mediator values.

The results are similar when we consider three-year mediator values for social bond strength with males (Figure S6). However, when we use three-year mediator values for social bond strength with females as the mediator, the mediated effect reached significance when at least one of the correlations was small (when $\zeta_M = 0$ and $\zeta_S \geq 0$, when $\zeta_M = -0.1$ and $\zeta_S \leq 0.5$, when $\zeta_M = -0.2$ and $\zeta_S \leq 0.1$, and when $\zeta_M \leq -0.5$ and $\zeta_S = 0$), but was not significant otherwise. All significant mediated effects were small and of similar magnitudes to the mediated effects estimated in our mediation analysis, suggesting the mediated effect through social bond strength with females is robust under a violation of sequential unconfoundedness. Thus, we still found a small, but significant mediated effect of cumulative adversity on survival via the path through bonds with females with modest violations of the sequential unconfoundedness assumption. This effect vanishes if violations to the assumption are strong.

### Supplementary Tables

**Table S1**. **Social bond strength with adult females as mediator, using one-year mediator values.** Mediation results from models that used one-year mediator values of social bond strength with adult females. Total, direct, mediated, and bond effect are measured in years. The effect on the mediator is measured in social bond strength units (i.e., DSI units; 1 SD in social bond strength with females=0.59 social bond strength units). Bolded effects are those for which the 95% credible interval did not overlap zero. Effect names are colored as in Figure 1.

| | Total effect ($\beta_2$, years) | Direct effect ($\beta_3$, years) | Mediated effect ($\beta_1\gamma$, years) | Effect on mediator ($\beta_1$, DSI units) | Bond effect ($\gamma$, years) |
|---|---|---|---|---|---|
| Drought | **-2.40 [-4.41, -0.39]** | **-2.03 [-3.63, -0.44]** | 0.36 [-0.09, 0.82] | **-0.21 [-0.38, -0.03]** | **1.80 [0.46, 3.13]** |
| Large group size | -1.59 [-4.14, 0.96] | -1.44 [-3.02, 0.14] | 0.15 [-0.01, 0.31] | -0.1 [-0.21, 0.01] | **1.70 [0.44, 2.96]** |
| Close-in-age younger sibling | -0.89 [-5.75, 3.97] | -0.69 [-2.12, 0.75] | 0.21 [-0.51, 0.92] | **-0.16 [-0.29, -0.03]** | **1.60 [0.52, 2.67]** |
| Maternal loss | **-3.20 [-5.61, -0.79]** | **-2.80 [-5.00, -0.59]** | 0.40 [-1.14, 1.94] | **-0.25 [-0.46, -0.04]** | **1.70 [0.52, 2.87]** |
| Low maternal social connectedness | 0.20 [-2.07, 2.48] | 0.21 [-1.31, 1.74] | 0.01 [-0.31, 0.33] | -0.04 [-0.15, 0.06] | **1.60 [0.53, 2.66]** |
| Low maternal social status | -1.79 [-4.51, 0.93] | -1.46 [-3.10, 0.18] | 0.33 [-0.04, 0.70] | **-0.15 [-0.26, -0.03]** | **1.70 [0.45, 2.94]** |
| Cumulative adversity | **-1.50 [-2.65, -0.34]** | **-1.36 [-2.39, -0.33]** | **0.14 [0.01, 0.26]** | **-0.09 [-0.16, -0.01]** | **1.79 [0.60, 2.98]** |

**Table S2. Social bond strength with adult males as mediator, using one-year mediator values.** Mediation results that used one-year mediator values for social bond strength with adult males. Total, direct, mediated, and bond effect are measured in years. The effect on the mediator is measured in social bond strength units (i.e., DSI units; 1 SD in social bond strength with males=0.70 units). Bolded effects are those for which the 95% credible interval did not overlap zero. Effect names are colored as in Figure 1.

| | Total effect ($\beta_2$, years) | Direct effect ($\beta_3$, years) | Mediated effect ($\beta_1\gamma$, years) | Effect on mediator ($\beta_1$, DSI units) | Bond effect ($\gamma$, years) |
|---|---|---|---|---|---|
| Drought | **-2.50 [-4.59, -0.40]** | **-2.17 [-3.88, -0.47]** | 0.32 [-0.04, 0.68] | **-0.15 [-0.28, -0.02]** | **2.10 [0.54, 3.66]** |
| Large group size | -1.49 [-3.9, 0.91] | -1.43 [-2.99, 0.14] | 0.07 [-3.09, 3.23] | -0.05 [-0.93, 0.83] | **2.10 [0.54, 3.66]** |
| Close-in-age younger sibling | -0.90 [-5.42, 3.63] | -0.72 [-2.13, 0.68] | 0.17 [-1.64, 1.98] | -0.11 [-0.61, 0.39] | **2.00 [0.65, 3.34]** |
| Maternal loss | **-3.30 [-5.78, -0.81]** | **-3.21 [-5.74, -0.68]** | 0.09 [-6.77, 6.95] | -0.06 [-1.95, 1.84] | **2.10 [0.64, 3.55]** |
| Low maternal social connectedness | 0.10 [-2.01, 2.21] | 0.33 [-1.29, 1.96] | 0.23 [-0.41, 0.87] | -0.15 [-0.32, 0.03] | **2.00 [0.66, 3.33]** |
| Low maternal social status | -1.80 [-4.42, 0.83] | -1.25 [-2.88, 0.38] | 0.55 [-0.06, 1.16] | **-0.24 [-0.43, -0.05]** | **1.90 [0.50, 3.29]** |
| Cumulative adversity | **-1.50 [-2.65, -0.34]** | **-1.38 [-2.43, -0.34]** | 0.12 [-0.94, 1.17] | -0.08 [-0.37, 0.22] | **1.80 [0.61, 2.99]** |

**Table S3. Moderation analyses, using one-year moderator values.** Moderation results for the effects of one-year social bond strength with females, social bond strength with males, and female social status. The bolded effects (those for which the 95% credible interval did not overlap zero) show that the effects of maternal loss on survival were moderated by social bond strength with both sexes and that the effects of low maternal social connectedness were moderated by adult social bond strength with males. Interaction effects are measured in terms of the log hazard ratio (log HR). Negative values indicate that higher values of the moderator (adult social bond strength or adult social status) act as a buffer, reducing the negative effects of early adversity on survival, while lower value of the moderator act as an amplifier, increasing the negative effects of early adversity on survival. A positive value indicates that higher values of the moderator act as an amplifier, while lower values of the moderator act as a buffer.

| | Social Bonds Females (log HR) | Social Bonds Males (log HR) | Social Status (log HR) |
|---|---|---|---|
| Drought | -0.15 [-0.37, 0.07] | -0.10 [-0.30, 0.09] | -0.08 [-0.35, 0.19] |
| Large group size | 0.15 [-0.10, 0.40] | -0.10 [-0.24, 0.05] | -0.14 [-0.33, 0.05] |
| Close-in-age younger sibling | 0.10 [-0.41, 0.60] | 0.00 [-0.33, 0.33] | 0.05 [-0.25, 0.35] |
| Maternal loss | **-0.20 [-0.33, -0.07]** | **-0.16 [-0.25, -0.07]** | 0.01 [-0.11, 0.14] |
| Low maternal social connectedness | 0.09 [-0.06, 0.23] | **-0.11 [-0.20, -0.03]** | -0.09 [-0.21, 0.03] |
| Low maternal social status | 0.01 [-0.15, 0.18] | 0.03 [-0.05, 0.12] | -0.16 [-0.44, 0.12] |
| Cumulative adversity | -0.02 [-0.04, 0.01] | -0.02 [-0.04, 0.00] | 0.03 [-0.01, 0.06] |

**Table S4. Incorporating hybrid score into mediation analysis for social bond strength with females.** Mediation results from models that used three-year mediator values of social bond strength with females, including hybrid score as a covariate. Total, direct, mediated and bond effects are measured in years. The effect on the mediator is measured in social bond strength units (i.e., DSI units). Bolded effects are those where the 95% credible interval did not overlap zero. Results are largely consistent with three-year mediator models (Table 2) that did not include hybrid score as a covariate, with similar effect sizes and confidence intervals. Effect names are colored as in Figure 1.

| | Total effect ($\beta_2$, years) | Direct effect ($\beta_3$, years) | Mediated effect ($\beta_1\gamma$, years) | Effect on mediator ($\beta_1$, DSI units) | Bond effect ($\gamma$, years) |
|---|---|---|---|---|---|
| Drought | **-2.69 [-4.95, -0.44]** | **-2.25 [-4.02, -0.48]** | **0.44 [0.03, 0.85]** | **-0.21 [-0.38, -0.03]** | **2.19 [0.56, 3.82]** |
| Large group size | -1.60 [-5.37, 2.17] | -1.38 [-3.44, 0.67] | 0.22 [-0.01, 0.44] | -0.11 [-0.26, 0.04] | **2.39 [0.61, 4.17]** |
| Close-in-age younger sibling | -0.90 [-6.55, 4.75] | -0.58 [-2.82, 1.66] | 0.32 [-0.04, 0.68] | **-0.16 [-0.28, -0.03]** | **2.3 [0.75, 3.85]** |
| Maternal loss | **-3.30 [-5.78, -0.81]** | **-2.67 [-4.77, -0.57]** | 0.63 [-0.22, 1.47] | **-0.26 [-0.48, -0.04]** | **2.59 [0.79, 4.39]** |
| Low maternal social connectedness | 0.11 [-2.96, 3.18] | 0.16 [-1.93, 2.26] | 0.05 [-0.09, 0.19] | -0.04 [-0.12, 0.04] | **2.59 [0.86, 4.33]** |
| Low maternal social status | -1.79 [-5.14, 1.56] | -1.35 [-3.54, 0.85] | 0.45 [-0.05, 0.94] | **-0.14 [-0.25, -0.03]** | **2.50 [0.66, 4.33]** |
| Cumulative adversity | **-1.59 [-2.82, -0.36]** | **-1.42 [-2.49, -0.34]** | **0.17 [0.01, 0.33]** | **-0.08 [-0.16, -0.01]** | **2.19 [0.74, 3.64]** |

**Table S5. Incorporating hybrid score into mediation analysis for social bond strength with males**. Mediation results from models that used three-year mediator values of social bond strength with males, including hybrid score as a covariate. Total, direct, mediated and bond effect are measured in years. The effect on the mediator is measured in social bond strength units. Bolded effects are those where the 95% credible interval did not overlap zero. Results are consistent with three-year mediator models (Table 3) that did not include hybrid score as a covariate with similar effect sizes and confidence intervals. Effect names are colored as in Figure 1.

| | Total effect ($\beta_2$, years) | Direct effect ($\beta_3$, years) | Mediated effect ($\beta_1\gamma$, years) | Effect on mediator ($\beta_1$, DSI units) | Bond effect ($\gamma$, years) |
|---|---|---|---|---|---|
| Drought | **-2.70 [-4.96, -0.44]** | **-2.34 [-4.18, -0.50]** | 0.36 [-0.05, 0.76] | **-0.16 [-0.29, -0.02]** | **2.39 [0.61, 4.17]** |
| Large group size | -1.60 [-5.24, 2.05] | -1.53 [-3.77, 0.72] | 0.07 [-0.19, 0.34] | -0.05 [-0.10, 0.00] | **2.40 [0.61, 4.18]** |
| Close-in-age younger sibling | -0.90 [-6.43, 4.64] | -0.70 [-2.68, 1.27] | 0.19 [-0.64, 1.03] | -0.11 [-0.52, 0.30] | **2.29 [0.75, 3.84]** |
| Maternal loss | **-3.29 [-5.78, -0.81]** | **-3.21 [-5.73, -0.68]** | 0.09 [-0.22, 0.40] | -0.05 [-0.11, 0.01] | **2.19 [0.67, 3.72]** |
| Low maternal social connectedness | 0.10 [-2.92, 3.12] | 0.36 [-1.93, 2.65] | 0.26 [-0.09, 0.60] | -0.14 [-0.30, 0.01] | **2.20 [0.73, 3.66]** |
| Low maternal social status | -1.79 [-5.19, 1.60] | -1.15 [-3.28, 0.99] | 0.65 [-0.07, 1.36] | **-0.24 [-0.43, -0.05]** | **2.30 [0.61, 3.99]** |
| Cumulative adversity | **-1.60 [-2.83, -0.36]** | **-1.45 [-2.55, -0.35]** | 0.15 [-0.09, 0.38] | -0.08 [-0.17, 0.01] | **2.19 [0.74, 3.65]** |

**Table S6. Adult female social status as mediator, using three-year mediator values**. Mediation results from models where three-year female social status was the mediator. Total, direct, mediated and status effect are measured in years. The effect on the mediator is measured in proportional rank units. Bolded effects are those where the 95% credible interval did not overlap zero. We see no effect of female social status on survival ('Status effect' has no bolded rows), no mediating effects of social status on the relationship between early adversity and survival ('Mediated effect' has no bolded rows), and among the sources of early adversity only maternal social status affects female social status (see 'Effect on mediator' column). Effect names are colored as in Figure 1.

| | Total effect ($\beta_2$, years) | Direct effect ($\beta_3$, years) | Mediated effect ($\beta_1\gamma$, years) | Effect on mediator ($\beta_1$, prop rank) | Status effect ($\gamma$, years) |
|---|---|---|---|---|---|
| Drought | **-2.60 [-4.77, -0.42]** | **-2.47 [-4.42, -0.53]** | -0.12 [-0.27, 0.03] | -0.14 [-0.31, 0.02] | 0.84 [-0.62, 2.29] |
| Large group size | -1.60 [-4.01, 0.81] | -1.73 [-3.62, 0.17] | 0.13 [-0.18, 0.44] | 0.14 [-0.01, 0.28] | 0.83 [-0.65, 2.31] |
| Close-in-age younger sibling | -0.89 [-5.55, 3.77] | -1.04 [-2.46, 0.38] | 0.15 [-0.02, 0.32] | 0.14 [-0.01, 0.29] | 0.89 [-0.53, 2.32] |
| Maternal loss | **-3.20 [-5.60, -0.79]** | **-3.18 [-5.69, -0.68]** | -0.01 [-0.14, 0.12] | -0.04 [-0.16, 0.09] | 0.85 [-0.61, 2.30] |
| Low maternal social connectedness | 0.11 [-1.85, 2.06] | 0.09 [-1.38, 1.56] | 0.02 [-0.18, 0.21] | -0.04 [-0.17, 0.10] | 0.79 [-0.69, 2.26] |
| Low maternal social status | -1.80 [-4.10, 0.51] | -0.79 [-2.60, 1.02] | -1.01 [-2.49, 0.48] | **-1.09 [-1.94, -0.24]** | 0.83 [-0.98, 2.64] |
| Cumulative adversity | **-1.60 [-2.83, -0.36]** | **-1.47 [-2.58, -0.36]** | -0.13 [-0.51, 0.25] | -0.21 [-0.46, 0.03] | 0.63 [-0.89, 2.16] |

### Supplementary figures

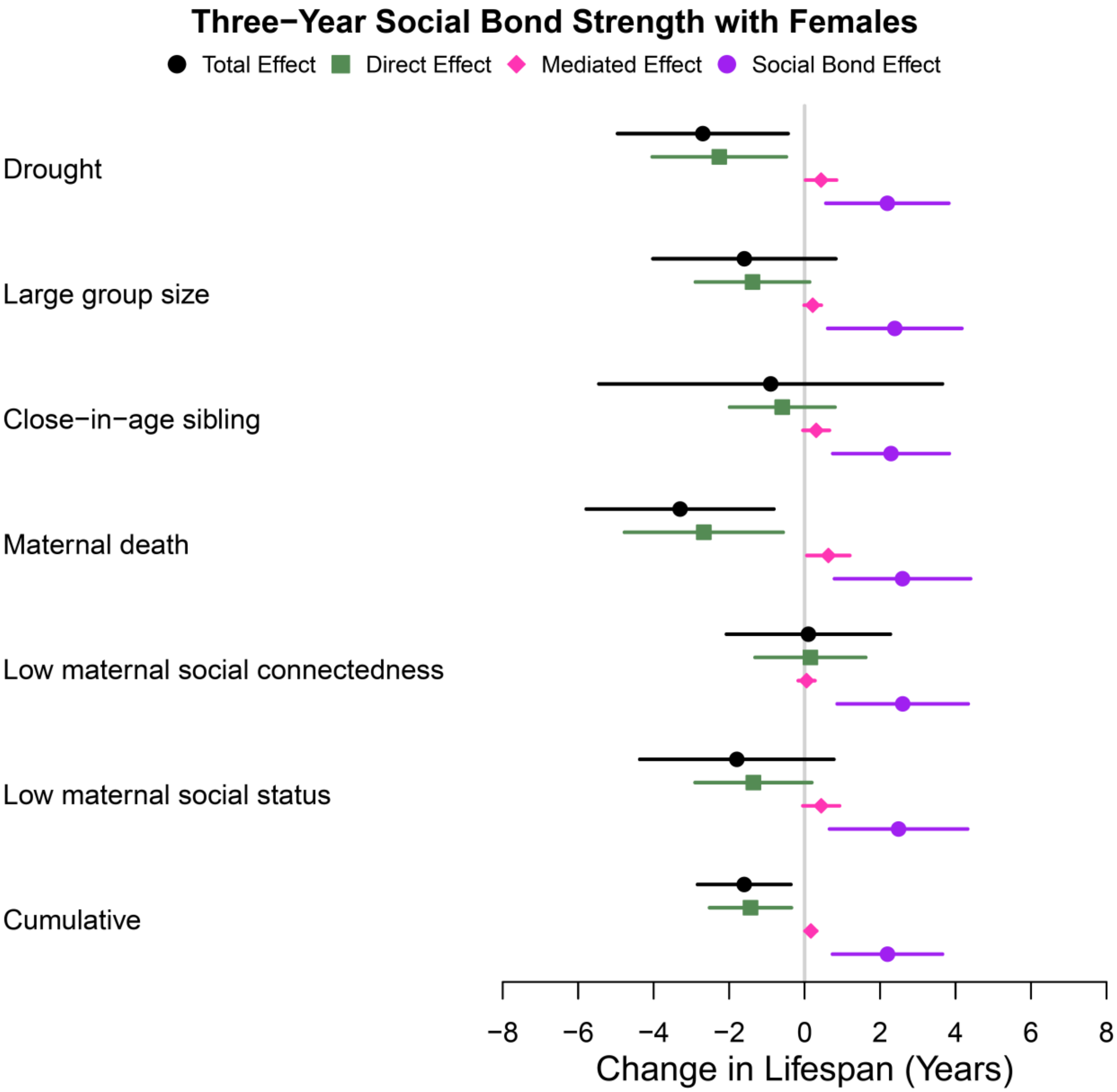


**Fig. S1.** Effects on survival, in models in which social bond strength with other females (estimated over three-year periods) was the mediator, showing effect sizes and 95% credible intervals for the total, direct, mediated, and social bond effects on survival.

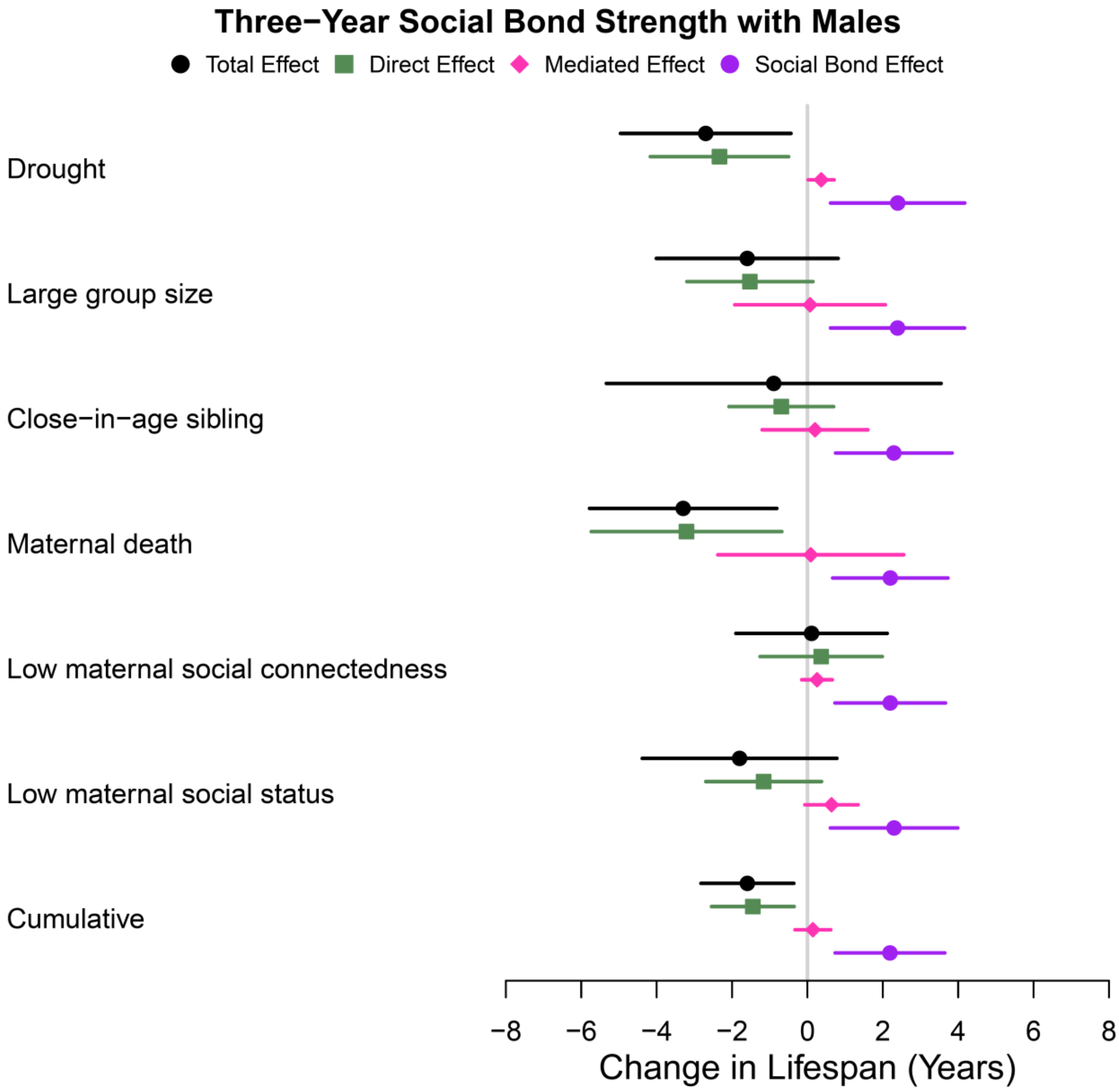


**Fig. S2.** Effects on survival, in models in which social bond strength with adult males (estimated over three-year periods) was the mediator, showing effect sizes and 95% credible intervals for the total, direct, mediated, and social bond effects on survival.

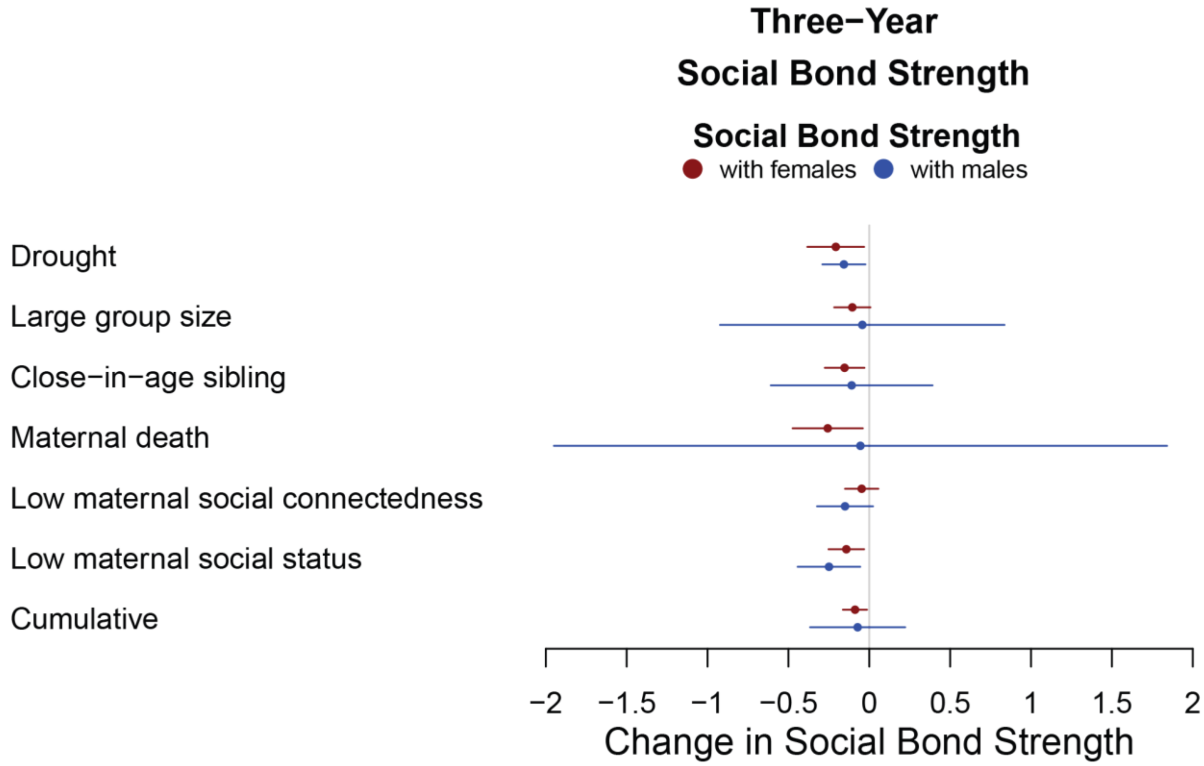


**Fig. S3.** Effects of early adversity on social bond strength, in models in which social bond strength with other adults of either sex (estimated over three-year periods) was the mediator, showing the effect sizes and 95% credible intervals for the relationship between early adversity and three-year social bond strength with females (red) or males (blue).

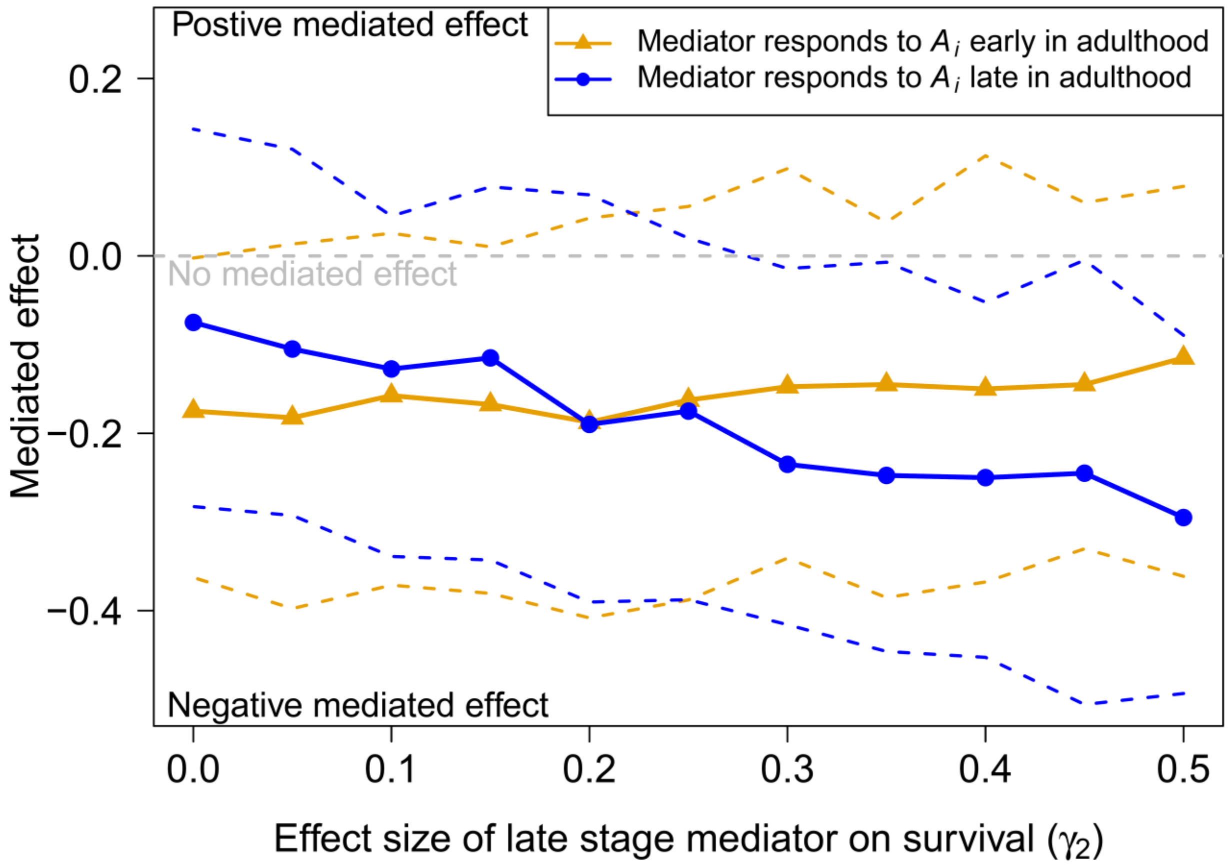


**Fig. S4.** Results of simulations showing that the strength of the mediated effect (y-axis) depends on the match or mismatch between timing of the effect of early adversity on the mediator (line colors; solid lines are the mean mediated effect sizes with dashed lines showing the 95% credible intervals.) and the timing of the effect of the mediator on survival (x-axis). The strongest mediated effects (more negative values on the y-axis) occur when timing is matched; i.e., when the effect of early adversity on the mediator occurs early in adulthood (orange line) and survival depends on the mediator early in adulthood (lower values of x-axis) or when the effect of early adversity on the mediator is greatest later in adulthood (blue line) and survival depends on the mediator value later in adulthood (higher values of the x-axis). More positive values of the y-axis correspond to a stronger mediated effect in which the pathway from early adversity to survival through the mediator increases survival. More negative values of the y-axis correspond to a stronger mediated effect in which the pathway from early adversity to survival through the mediator decreases survival. A value of zero on the y-axis indicates no mediated effect. Larger values of the x-axis correspond to simulations in which the effect of the mediator at the later stage (eqn. S1b) on survival later in life (stage 2) was stronger than the effect of the mediator at the early stage (stage 1) on survival to stage 2 (e.g, $\gamma_1 < \gamma_2$; the effect of the mediator later in adulthood affected survival late in life). Smaller values of the x-axis correspond to simulations in which the effect of the mediator at the early stage (stage 1; Equation S1a) on survival to stage 2 was stronger than the effect of the mediator at the later stage (Equation S1b) on survival to stage 2 (e.g., $\gamma_1 > \gamma_2$; the effect of the mediator earlier in life affected survival later in adulthood). The orange line corresponds to cases where the effect of early adversity on the mediator was strongest early in adulthood (Equation S1a), and the blue line corresponds to cases where the effect of early adversity on the mediator was strongest late in adulthood (Equation S1b). We found that even when we fixed the effect of early adversity on the mediator and the effect of the mediator on survival, the mediated effect size changes when the strength of these effects is time-varying.

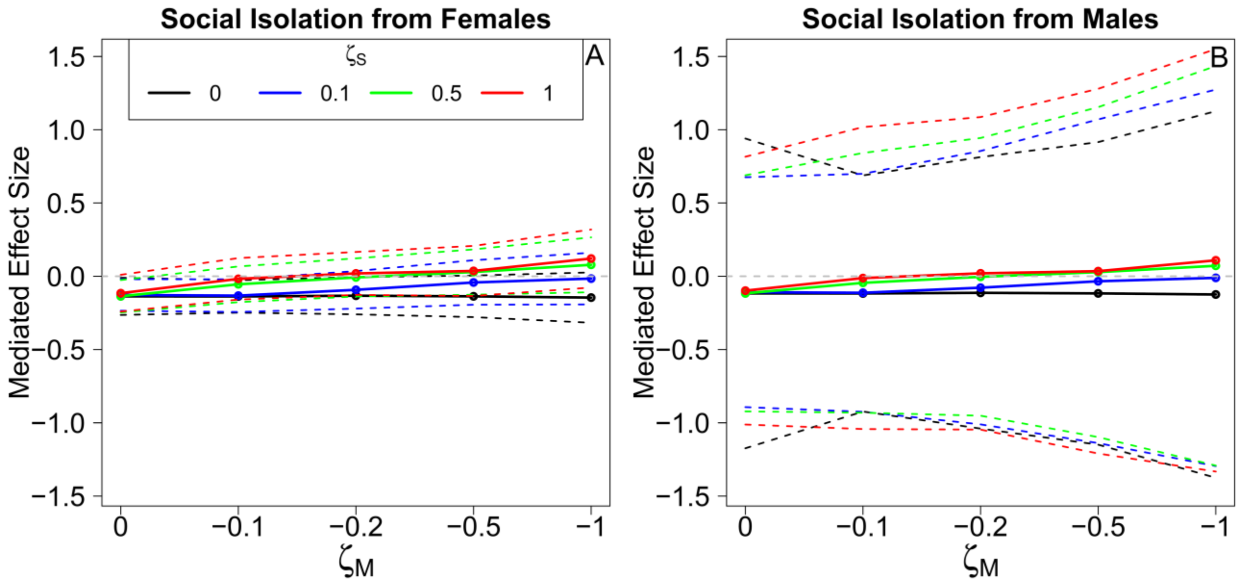


**Fig. S5.** Results of a sensitivity analysis to test whether model results are robust to violations of the assumption of sequential unconfoundedness: one-year mediator values were used in the analysis, see Fig. S6 for sensitivity analysis with three-year mediator values. (A) Varying the correlation between the unmeasured confounder and mediator, $\zeta_M$ (x-axis), and the unmeasured confounder and the survival outcome, $\zeta_S$ (line colors), does not change the small and largely not significant estimate of the mediated effect for models where the mediator was social bond strength with females. (B) The same is true for social bond strength with males. Therefore, we find no evidence that sequential unconfoundedness affects our estimates of the mediated effect for our one-year mediator models. Solid lines are the mean mediated effect sizes with dashed lines showing the 95% credible intervals.

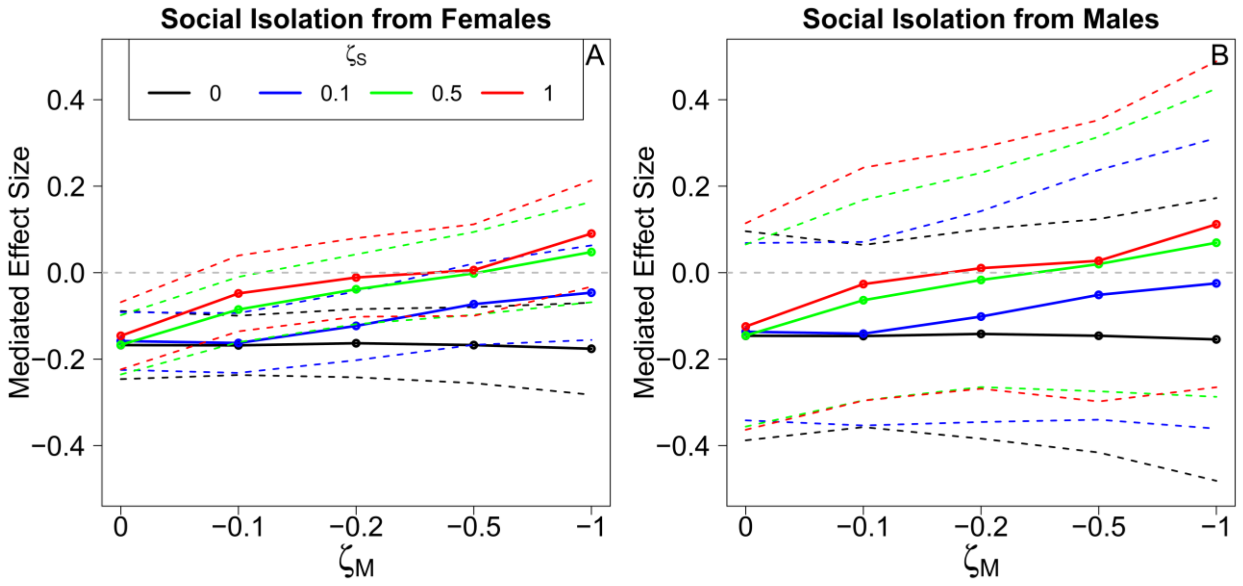


**Fig. S6.** Results of a sensitivity analysis to test whether model results are robust to violations of the assumption of sequential unconfoundedness; three-year mediator values were used in this analysis. (A) Varying the correlation between the unmeasured confounder and mediator, $\zeta_M$ (x-axis), and the unmeasured confounder and the survival outcome, $\zeta_S$ (line colors), significantly affected the mediated effect size when the mediator was three-year social bond strength with females for a range of values for $\zeta_M$ and $\zeta_S$ (when $\zeta_M = 0$ and $\zeta_S \geq 0$, when $\zeta_M = -0.1$ and $\zeta_S \leq 0.5$, when $\zeta_M = -0.2$ and $\zeta_S \leq 0.1$, and when $\zeta_M \leq -0.5$ and $\zeta_S = 0$). However, all significant mediated effects were small and of similar magnitudes to the mediated effects estimated in our mediation analysis, suggesting that even in these cases, the mediated effect through social bond strength with females is robust under a violation to sequential unconfoundedness. Furthermore, no effect was seen when correlations were stronger. (B) Varying the correlation between the unmeasured confounder and the mediator and the unmeasured confounder and survival does not change the small and non-significant estimate of the mediated effect for three-year social bond strength with males. Solid lines are the mean mediated effect sizes with dashed lines showing the 95% credible intervals.
